# Association of microglia loss with hippocampal network impairments as a turning point in the amyloid pathology progression

**DOI:** 10.1101/2024.03.14.584993

**Authors:** Giusy Pizzirusso, Efthalia Preka, Julen Goikolea, Celia Aguilar-Ruiz, Patricia Rodriguez Rodriguez, Guillermo Vazquez Cabrera, Simona Laterza, Maria Latorre Leal, Francesca Eroli, Klas Blomgren, Silvia Maioli, Per Nilsson, Adamantia Fragkopoulou, André Fisahn, Luis Enrique Arroyo-García

## Abstract

Alzheimer’s disease is a progressive neurological disorder causing memory loss and cognitive decline. The underlying causes of cognitive deterioration and neurodegeneration remain unclear, leading to a lack of effective strategies to prevent dementia. Recent evidence highlights the role of neuroinflammation, particularly involving microglia, in Alzheimer’s disease onset and progression. Characterizing the initial phase of Alzheimer’s disease can lead to the discovery of new biomarkers and therapeutic targets, facilitating timely interventions for effective treatments. We used the *App^NL-G-F^* knock-in mouse model, which resembles the amyloid pathology and neuroinflammatory characteristics of Alzheimer’s disease, to investigate the transition from a pre-plaque to an early plaque stage with a combined functional and molecular approach. Our experiments show a progressive decrease in the power of cognition-relevant hippocampal gamma oscillations during the early stage of amyloid pathology, together with a modification of fast-spiking interneuron intrinsic properties and postsynaptic input. Consistently, transcriptomic analyses revealed that these effects are accompanied by changes in synaptic function-associated pathways. Concurrently, homeostasis-and inflammatory-related microglia signature genes were downregulated. Moreover, we found a decrease in Iba1-positive microglia in the hippocampus that correlates with plaque aggregation and neuronal dysfunction. Collectively, these findings support the hypothesis that microglia play a protective role during the early stages of amyloid pathology by preventing plaque aggregation, supporting neuronal homeostasis, and overall preserving the oscillatory network’s functionality. These results suggest that the early loss of microglia could be a pivotal event in the progression of Alzheimer’s disease, potentially triggering plaque deposition, impairment of fast-spiking interneurons, and the breakdown of the oscillatory circuitry in the hippocampus.

## 1 Introduction

Alzheimer’s disease (AD) is a chronic multifactorial disorder characterized by progressive memory loss and cognitive impairment, preceded by a decades-long asymptomatic stage [1]. The events that drive the onset of the pathology leading to cognitive decline and neurodegeneration in the advanced stages remain unclear. This poor understanding of AD aetiopathogenesis is reflected in the absence of early treatment strategies to prevent the onset of dementia rather than slowing down its progression [2,3]. Amyloid beta (Aβ) accumulation is traditionally considered a major AD driver [4,5], however, growing evidence suggests that Aβ amyloidosis alone is not enough to explain the entire AD pathogenesis [6]. Among others, neuroinflammation has been proposed as an important contributor to AD onset and progression [7,8]. Indeed, epidemiological studies have reported a positive association between cognitive decline and previous inflammatory events [9,10]

For this reason, microglia, as the resident immune cell type in the brain, are receiving increasing attention in AD studies. However, whether microglia play a protective or detrimental role during AD progression remains controversial [7,11]. The beneficial role of microglia in containing plaque deposition and delaying cognitive impairment has been reported by several studies [12–16]. Nonetheless, chronic activation of microglia is a well-known cause of neuroinflammation and neurodegeneration [17–19].

Besides their immune-related functions, microglia are involved in homeostatic processes in the brain, including neuronal and network activity regulation [20–23]. One of the consequences of AD is the dysfunction of multiple brain networks, resulting in the inability of the circuits to synchronize in specific oscillation rhythms [24,25]. In particular, brain oscillations in the gamma-frequency band (30-80 Hz) play a pivotal role in higher brain activities like information processing and memory formation [26–28]. We previously reported that disruption of cognition-relevant gamma oscillations and fast-spiking interneurons (FSI) rhythmicity occurs at an early stage of amyloid pathology in the hippocampus of the *App^NL-G-F^* mouse model [29]. This functional impairment takes place before widespread plaque deposition in the hippocampus. Importantly, restoration of FSI rhythmicity and hippocampal oscillatory activity leads to improved cognitive performance [30,31], underlining the importance of hippocampal network dysfunction in AD pathology progression.

Here, we used a combination of functional and molecular approaches to investigate the mechanisms behind the failure of the hippocampal network and the role of microglia at the earliest stages of amyloid pathology. To our knowledge, this is the first time that such an early pre-plaque stage of amyloidosis progression has been investigated by RNA sequencing in this model. Elucidating the events characterizing the initial phase of the amyloid pathology can generate new perspectives on AD’s etiopathogenesis, hence providing new biomarkers and targets for the development of a therapeutic approach to AD based on timely intervention.

## 2 Materials and methods

### 2.1 Animals

Experiments were performed according to the European and Swedish animal welfare regulations (ethical permit: N45/13 and 12570-2021 approved by Stockholm Animal Ethical Board). Male homozygous *App^NL-G-F^* [32] and wild-type (WT) littermates were bred using the C57/BL6J strain as a background. The mice were sacrificed for brain collection by decapitation under deep anesthetization with isoflurane. Brains were collected between post-natal day (P) 30 and 65 and divided into four representative age points: P30 (P30-35), P40 (P36-45), P50 (P46-55) and P60 (P56-65).

### 2.2 Ex-vivo electrophysiological recordings

Following brain dissection, 350µm thick horizontal hippocampal sections were obtained and kept alive for electrophysiology recordings as previously reported [29]. All electrophysiological experiments were performed at a constant temperature of 34°C in a submerged-type recording chamber where slices received a continuous supply of 3 ml/min oxygenated artificial cerebrospinal fluid (ACSF) containing in mM: 124 NaCl, 30 NaHCO3, 10 glucose, 1.25 NaH2PO4, 3.5 KCl, 1.5MgCl2, 1.5 CaCl2. Recordings were performed using a Multiclamp 700B amplifier and acquired using pCLAMP 10,4 software (Molecular Devices). Signals were low pass filtered at 1 kHz, acquired at 5 kHz, digitized, and stored using Digidata 1322A and pCLAMP 10,4 software (Molecular Devices, CA, USA).

### 2.3 Ex vivo Gamma oscillations

For Local field potential (LFP) measurements, brains from all age points were used: P30 (WT n=6, *App^NL-G-F^* n=6), P40 (WT n=7, *App^NL-G-F^* n=6), P50 (WT n=6, *App^NL-G-F^* n=9), P60 (WT n=7, *App^NL-G-F^* n=11). Ex vivo gamma oscillations were induced by adding kainic acid (KA; Tocris Bioscience, Bristol, UK) at a concentration of 100 nM to the extracellular bath and were recorded using borosilicate glass microelectrodes (1,5-2,5 MΩ) filled with ACSF placed in the stratum oriens of the CA3 area of the hippocampus. The oscillations were allowed to stabilize for at least 20 minutes before recordings were performed. Signals were conditioned using a HumBug 50 Hz noise eliminator (Quest Scientific). For oscillation power spectra Fast Fourier Transformations were obtained from 60 s of LFP recording (segment length 8192 points) using Axograph software (Kagi, Berkeley, CA, USA). Frequency variance data was obtained from the power spectra described above using Axograph X. Gamma power was calculated by integrating the power spectral density from 20 to 80 Hz using Clampfit 11.2.

### 2.4 Intrinsic properties of fast-spiking interneurons

Intrinsic properties and spontaneous excitatory post-synaptic currents (sEPSC) were recorded in whole-cell patch clamp mode from fast-spiking interneurons (FSI) in the CA3 stratum radiatum of hippocampal slices from P30 (WT n=17, *App^NL-G-F^* n=10) and P60 (WT n=11, *App^NL-G-F^* n=17) brains. Patch clamp recordings were performed with borosilicate glass micropipettes (4–6 MΩ) filled with a potassium-based internal recording solution (in mM: 122.5 K+-gluconate, 8 KCl, 2 Mg2+ ATP, 0,3 Na+ GTP, 10 HEPES, 0,2 EGTA, 2 MgCl; pH 7.2–7.3; osmolarity 270–280 mosmol/l) containing 2% of neurobiotin (Vector Laboratories) as a neurotracer. FSI were visualized under an upright microscope using IR-DIC microscopy (Axioskop, Carl Zeis AG, Göttingen, Germany) and identified based on their location, morphology, and response to custom-made current delivery protocols as described previously [29,33]. EPSCs were recorded in voltage-clamp configuration with the voltage holding at -70 mV. Firing and membrane potential properties were recorded in gap-free current-clamp mode. Each cell classified as FSI underwent consequent hyper-and de-polarizing current steps (Input resistance protocol [34]: 25 steps of 10pA from -100pA to +140pA, 700ms each; Fig 2.A) starting from the membrane resting potential and from a -70mV membrane potential. After recording, the slice was stored in PFA overnight and then in a 30% sucrose solution until further experiments. All recordings obtained from FSI were analyzed using a costume-made Python script. sEPSC were analyzed using the Pyabf and Scipy packages. Firing and membrane intrinsic properties were obtained from the abovementioned input resistance protocol using the Pyabf, Ipfx, and Efel packages.

- Rehobase (pA), firing threshold (mV), and firing latency (ms) were extracted from the first sweep of the input resistance protocol with at least one action potential.
- Mean action potential amplitude (mV), half-width (ms), peak upstroke (V/s), peak downstroke (V/s), rise rate, fall rate, rise time (ms), and fall time (ms) were obtained from the first sweep of the input resistance protocol with four to six action potentials.
- Firing rate (Hz) was calculated as the number of action potentials fired during the input resistance protocol, divided by the duration of the protocol.
- Ohmic input resistance (GΩ) was calculated by averaging the ohmic input resistance values from subthreshold hyper-and depolarizing current steps in the input resistance protocol (from -30 to +30 mV) in the absence of action potentials.
- Membrane potential at the corresponding current step was extracted from each trace of input resistance protocol. Resting membrane potential (Em0) corresponds to the membrane potential when no current is injected.
- Sag (mV) was calculated from the most hyperpolarized sweep (-100 pA) of the input resistance protocol recorded with the voltage holding at -70mV.

### 2.5 RT-qPCR

WT (P30 n=4) and *App^NL-G-F^* (P30 n=4, P60 n=4) hippocampal slices were placed in a submerged recording chamber where a solution of ACSF and KA (100 nM) was constantly supplied for 2 hours. After 2 hours, gamma oscillations were recorded to confirm the network activation. For the control group, hippocampal slices from each group were incubated for 2 hours in ACSF without KA.

RNA was purified using the PureLink RNA mini kit with in-column DNAse-I digest (#79254, Qiagen) following the manufacturer’s instructions (#12183018A, ThermoFisher). RNA was then quantified in a nanodrop and reverse-transcribed using the SuperScript III First-Strand Synthesis System (#18080051, ThermoFisher), followed by RNAse H digestion. cDNA was then used to run qPCRs using TaqMan Universal PCR master mix.

FAM-labeled TaqMan assays (ThermoFisher) used were: Mm00463644_m1 (*Npas4*), and Mm00487425_m1 (*Fos*). *Gapdh* was used as endogenous control (4352932E). All qPCRs were performed in a 7500 Fast Real-Time PCR system.

### 2.6 RNA extraction and bulk RNA sequencing

Sixteen hippocampi (WT P30 n=4, WT P60 n=4, *App^NL-G-F^* P30 n=4, *App^NL-G-F^* P60 n=4) were dissected and used for bulk RNA-seq. Total RNA was isolated using the RNeasy Plus mini kit (QIAGEN #74134) according to the manufacturer’s instructions and processed for sequencing at the Bioinformatics and Expression Analysis core facility at Karolinska Institutet. Following the quality control by Agilent Bioanalyzer 2100, samples were processed for library preparation and sequenced using the Illumina Nextseq 2000 P3 (100 cycles) system. All the following analyses were performed in R (version 4.3.0). The DESeq2 (version 1.40,2) package was used to identify differentially expressed genes across different groups. Log fold changes obtained with a Wald test were shrunk using the apelgm method. Statistical significance was considered when adjusted p-value <0,05, as calculated with the Benjamini & Hochberg multiple adjustment method. For the functional classification of DEGs, we used the clusterprofiler [35,36] tool (version 4.8.2). Specifically, Gene Ontology (GO) enrichment analysis was performed to identify the activated and suppressed GO terms (biological processes, molecular function, and cellular component). GO terms that displayed a p-value <0,05, as derived by the hypergeometric test were sorted out as statistically significant. Principal component analysis (PCA) on the sequencing data was performed using the VSN package following instructions from the Bioconductor repository. Centroid coordinates for the PCA plot were calculated as the average of the x and y coordinates of individual samples belonging to the same group.

### 2.7 Western blot

Mouse hippocampal tissue was disrupted and homogenized in 1xRIPA buffer (#89901, Thermo Fisher Scientific) containing phosphatase and protease inhibitors (#P8340 and #P0044, Sigma-Aldrich). Homogenates were then centrifuged at 15000×g for 15 min 4 °C, and supernatants were mixed with 4X Protein Sample Loading Buffer (#928-40004, LI-COR) containing 2% β-mercaptoethanol (#444203, Sigma-Aldrich). Processed samples were then run in 12.5% homemade polyacrylamide gels (Bio-Rad) and transferred to a 0.45 NC nitrocellulose membranes (Amersham Protran 0.45 NC nitrocellulose membrane, # 10600002, VWR) for 2 h room temperature at 50 V. Membranes were then blocked for 30 minutes in 5% skim milk in 0.1% TBS-Tween-20 (TBS-T) prior to overnight incubation with diluted primary antibody at 4 °C. Primary antibodies: 1:1000 anti-GABA A Receptor alpha 1 polyclonal rabbit antibody (#ab33299, Abcam), 1:1000 anti-Glutamate Receptor 1 (AMPA subtype) polyclonal rabbit antibody (#ab31232, Abcam), 1:10000 anti-GAPDH monoclonal rabbit antibody (#ab8245, Abcam), 1:1000 anti-Iba1 polyclonal rabbit antibody (#ab153696, Abcam), 1:1000 anti-HER4/ErbB4 (111B2) Rabbit monoclonal antibody (#4795, Cell Signaling Technology), 1:1000 anti-KCNS3 polyclonal rabbit antibody (#PA5-96856, Invitrogen), 1:1000 anti-Nav1.1 Na+ Channel monoclonal mouse antibody (#75-023, Antibodiesinc), 1:1000 anti-KV3.1 (KCNC1) polyclonal rabbit antibody (#PA5-101763, Invitrogen), 1:1000 anti-Synaptotagmin-2 polyclonal rabbit antibody (#60889, Cell Signaling Technology), 1:1000 anti-GABA-A Receptor beta 3 polyclonal rabbit antibody (#NB300-199, Novus Biologocals, Biotechne), 1:500 anti-Glutamate receptor AMPA R4 (#G5790, Sigma-Aldrich). Fluorescent secondary antibodies (1:10000, LI-COR Biosciences) were used for 2 h at room temperature, and bands were visualized using ODYSSEY Infrared Imaging System (LI-COR Biosciences). Band intensity signal was quantified by ImageJ software. Each band signal value was normalized against loading control (GAPDH signal value).

### 2.8 Immunostaining and imaging

Thirty-seven (WT P30 n=11, WT P60 n=9, *App^NL-G-F^* P30 n=6, *App^NL-G-F^* P60 n=11) hippocampal slices containing neurobiotin-marked neurons were used for immunostaining. Non-specific binding was blocked by incubating the sections in a solution of 10% normal donkey serum (Jackson ImmunoResearch Laboratories, West Grove, PA), 1.5% Triton X-100 (made Tris buffer saline, TBS) for 2 h at room temperature. Sections were then incubated with primary antibodies rabbit anti-Iba1 (1:1000; WAKO) and mouse anti-human amyloid beta 82E1 (1:1000; IBL) at 4°C for 48 h and then for 2 h at room temperature with the fluorescent secondary antibodies Alexa-555 donkey anti-rabbit (1:1000; Abcam), Alexa-633 donkey anti-mouse (1:250; Biotium) and Streptavidin 488 (1:1000; Vector laboratories). Hoechst 33342 (Molecular Probes/Life Technologies) was used as a nuclear counterstain. Sections were mounted onto glass slides and coverslipped using ProLong diamond anti-fade reagent (Molecular probes/Life technologies). The sections were imaged using a confocal laser scanning microscope (Zeiss LSM900-Airyscan confocal) at the Biomedicum Imaging Core facility at Karolinska Institutet.

#### 2.8.1 Microglia and amyloid plaque density

Images of the entire hippocampus were acquired, using a 20x objective, by a combination of tiles and a Z-stack of a thickness of 10µm. These images were processed using the Zen system and their maximum intensity projection was obtained, which allowed for the quantification of the microglia using the Zeiss Zen 3.7 software. Furthermore, the area of the hippocampus was obtained using the lasso tool in the Zeiss software which was also used to acquire the area of the amyloid beta plaques. Microglia and amyloid plaque density as well as plaque area were calculated as a function of the hippocampal area.

#### 2.8.2 Microglia and FSI interaction

In twenty-five of these sections (WT P30 n=6, WT P60 n=6, *App^NL-G-F^* P30 n=6, *App^NL-G-F^* P60 n=6, containing one neurobiotin pre-filled neuron each) images of the neurobiotin-marked neurons were taken at a magnification of 0,7, using a 63x oil objective along with a Z-stack technique of a thickness of 10µm. Subsequently, these images underwent processing utilizing the Zen system: their maximum intensity projections, as well as a subset of images measuring 1355 x 1355 pixels, were generated. Cell surface creation was carried out using the IMARIS 10.0.1 software, by default parameters (surface detail 0.141 µm) in green (AF488) and red (AF555) channels. Surfaces were masked setting a constant in the voxel intensity. Outside surfaces intensity was set to value = 0 and inside surfaces to value =1, generating two new binary channels.

Using the IMARIS image processing tool “Channel Arithmetic” by Matlab XTension, the two newly generated binary channels were combined following the formula (chX * chY) resulting in a new channel (pink) that shows only the voxel overlaps from the two binary channels. Voxels with value 1 after the channel arithmetic correspond to the places where voxels of both channels had an overlap in positive signal, indicating the contact surface between cells. Image J software was used to obtain the area of the contacts between microglia and neurons and categorize the contact points into somatic and dendritic.

### 2.9 Statistical analysis

All statistical analyses, excluding the analysis of the bulk RNA sequencing data, were performed using GraphPad Prism 9.4.0. We performed two-sided statistical analyses comparing the *App^NL-G-F^* group and the WT at each age. We used a normality and lognormality test (Shapiro–Wilk test) and we excluded outliers after ROUT (Q = 1%) analysis. We performed one and two-way ANOVA followed by a Holm–Sidak’s multiple comparisons test, unpaired t-test, and Pearson correlation test according to the statistical purpose and number of groups in the analyses. In the graphs data are reported as means ± SEM and significance levels are *p < 0,05, **p < 0,01, ***p < 0,001, **** p < 0,0001. The three-dimensional PCA of neuronal properties was performed in Phyton using the Sklearn package.

## 3 Results

### 3.1 Progressive impairment of hippocampal gamma oscillations in *App^NL-G-F^* mice

Previous evidence from our lab shows an early impairment in the hippocampal oscillatory activity during amyloid pathology progression in the *App^NL-G-F^* knock-in mouse model [29]. We reported that gamma oscillations were strongly decreased in the *App^NL-G-F^* mice at P75 when compared to age-matched WT mice, while no differences were observed at P30 [29]. In the present study, we first aimed to narrow down the time-point when this shift in hippocampal gamma oscillations occurs. For this purpose, we investigated ex vivo gamma oscillations in hippocampal slices of *App^NL-G-F^* and WT mice at P30, P40, P50, and P60 (Fig 1.A).

**Fig 1.**
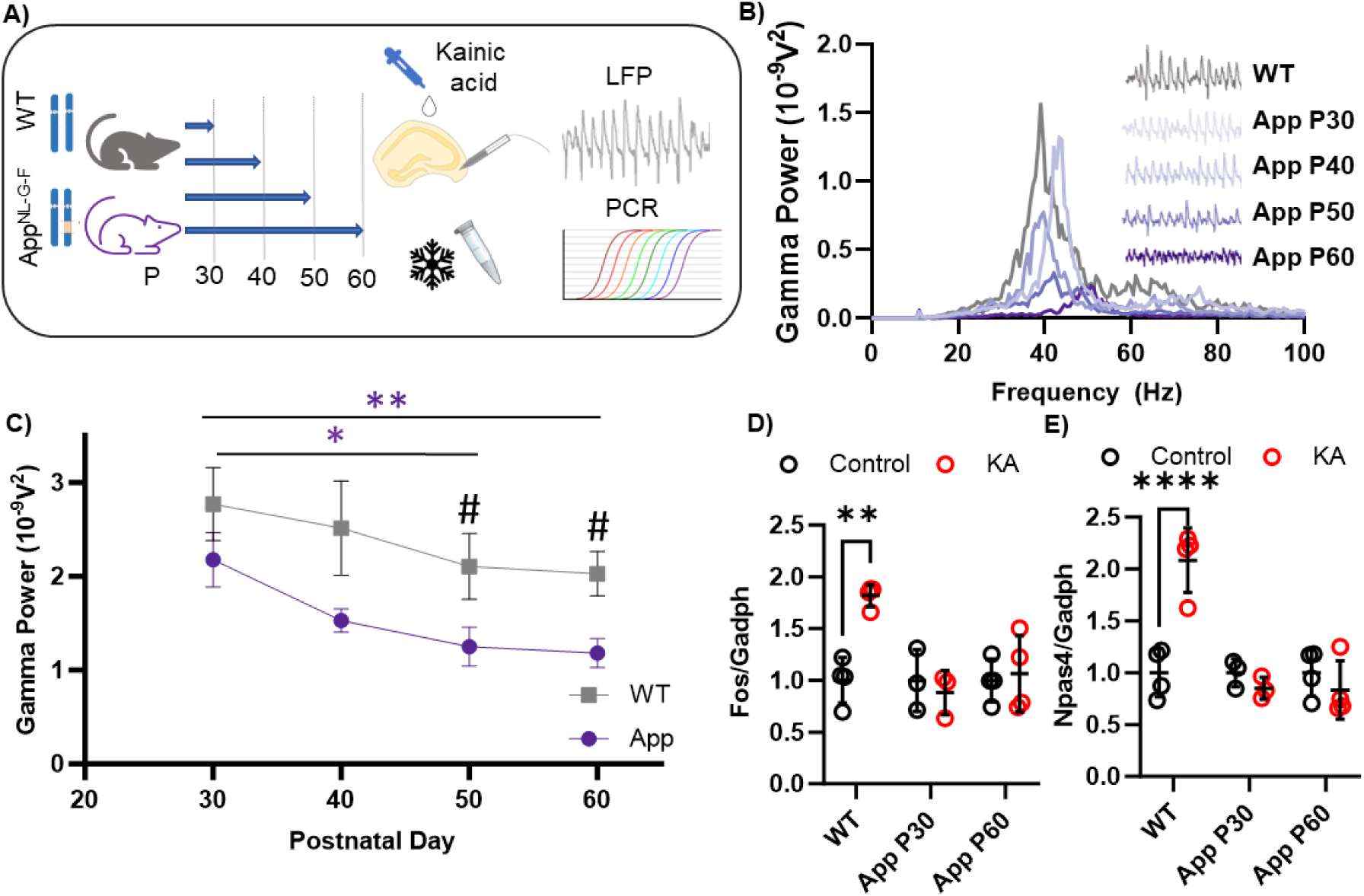
Progressive impairment of Hippocampal gamma oscillations in *App^NL-G-F^* mice. **A)** Graphic methods for LFP recordings and PCR. **B)** Representative power spectra from WT and *App^NL-G-F^* at different ages. **C)** Line plot of gamma oscillation power in WT and *App^NL-G-F^* mice at different ages showing a progressive decrease of the hippocampal network synchrony in *App^NL-G-F^* mice. The graph shows a two-way ANOVA with Holm–Sidak’s multiple comparisons test at P30 (WT n= 6 vs *App^NL-G-F^* n=6, p=0,124), P40 (WT n= 6 vs *App^NL-G-F^* n=6, p=0,172), P50 (WT n= 6 vs *App^NL-G-F^* n=9, p=0,017), and P60 (WT n= 7 vs *App^NL-G-F^* n=11, p=0,010) (# signficance). Intra-group differences were evaluated too: for the WT group P30 vs P40 (p= 0,287), P30 vs P50 (p= 0,304), P30 vs P60 (p=0,255). For the *App^NL-G-F^* group P30 vs P40 (p= 0,326), P30 vs P50 (p= 0,049), P30 vs P60 (p= 0,026) (* significance). **D)** Two-way ANOVA with Holm–Sidak’s multiple comparisons test for *Fos* normalized expression showing overexpression in WT KA-exposed slices (WT control n=4 vs KA n=4, p= 0,003) but not in *App^NL-G-F^* KA - exposed slices (*App^NL-G-F^* P30 control n=3 vs KA n=3, p=0,999; *App^NL-G-F^* ^F^ P60 n=4 control vs KA n=4, p=0,999). **E)** Two-way ANOVA with Holm–Sidak’s multiple comparisons test for *Npas4* normalized expression showing overexpression in WT KA-exposed slices (WT control n=4 vs KA n=4, p<0,0001) but not in *App^NL-G-F^* KA - exposed slices (*App^NL-G-F^* P30 control n=3 vs KA n=3, p=0,983; *App^NL-G-F^* P60 n=3 control vs KA n=3, p=0,983). Data are presented as mean ± SEM, and “n” indicates the number of mice. *p < 0,05, #p < 0,05, **p < 0,01, ***p < 0,001, **** p < 0,0001.

*App^NL-G-F^* mice exhibited a progressive decrease in gamma power that is significant at P50 compared to P30 mice (*App^NL-G-F^* P30 vs P50 p=0,027; Fig 1.B, C). Additionally, we confirmed that gamma oscillations were comparable in power comparing the *App^NL-G-F^* and WT groups at P30 (WT vs *App^NL-G-F^* P30 p=0,190; Fig 1.C), while they were significantly decreased at P50 in the *App^NL-G-F^* group as compared to age-matched WT mice (WT vs *App^NL-G-F^* P60 p=0,041; Fig 1.C). This gamma oscillation impairment in *App^NL-G-F^* mice continued to decrease at P60 (WT vs *App^NL-G-F^* P60 p=0,082, *App^NL-G-F^* P30 vs P60 p=0,014; Fig 1.C). No change was detected in frequency variance (Supp fig 1.A) and peak frequency (Supp fig 1.B).

These results provide a specific time window to investigate the functional and molecular changes in this early stage of amyloid pathology progression and potentially identify new targets to prevent and/or recover them. Therefore, we selected a time point in the *App^NL-G-F^* mice where the oscillatory activity is comparable to the WT mice (P30) and a time point with oscillatory impairment (P60) for further analysis.

After selecting the time points of interest, we next asked if the induction of hippocampal gamma oscillations by KA could promote the expression of immediate early genes (IEG). IEG are rapidly induced in response to neuronal depolarization and play an essential role in neuronal synaptic plasticity and learning-associated processes [37–40]. We therefore quantified *Fos* and *Npas4* gene expression analysis by RT-qPCR in KA-exposed hippocampal slices before and after gamma oscillation impairment in *App^NL-G-F^* mice and compared them to a WT group (Fig 1.A).

Our results show that the network activation with KA caused an induction of *Fos* (Fig 1.D, p=0,0005) and *Npas4* (Fig 1.E, p=0,0014) expression in WT mice. This effect was not observed in *App^NL-G-F^* mice at P30 nor at P60 (Fig 1.D, E), in line with previous studies that report suppression of IEG in the presence of Aβ [41].

### 3.2 Fast-spiking interneuron intrinsic properties and postsynaptic input change during amyloid pathology progression

Fast-spiking interneurons (FSI) are a subtype of GABAergic interneurons that hold a pivotal role in initiating and sustaining gamma oscillations [42–44]. We have previously reported that FSI spike-gamma coupling is affected during amyloid pathology progression at P75 [29]. Here we aimed to assess the functionality of FSI in the *App^NL-G-F^* mice. For this reason, we used the whole-cell patch-clamp technique to assess the intrinsic properties and post-synaptic currents of FSI in a resting state (without KA) in WT and *App^NL-G-F^* mice at P30 and P60 (Fig 2.A; Table 1).

**Fig 2.**
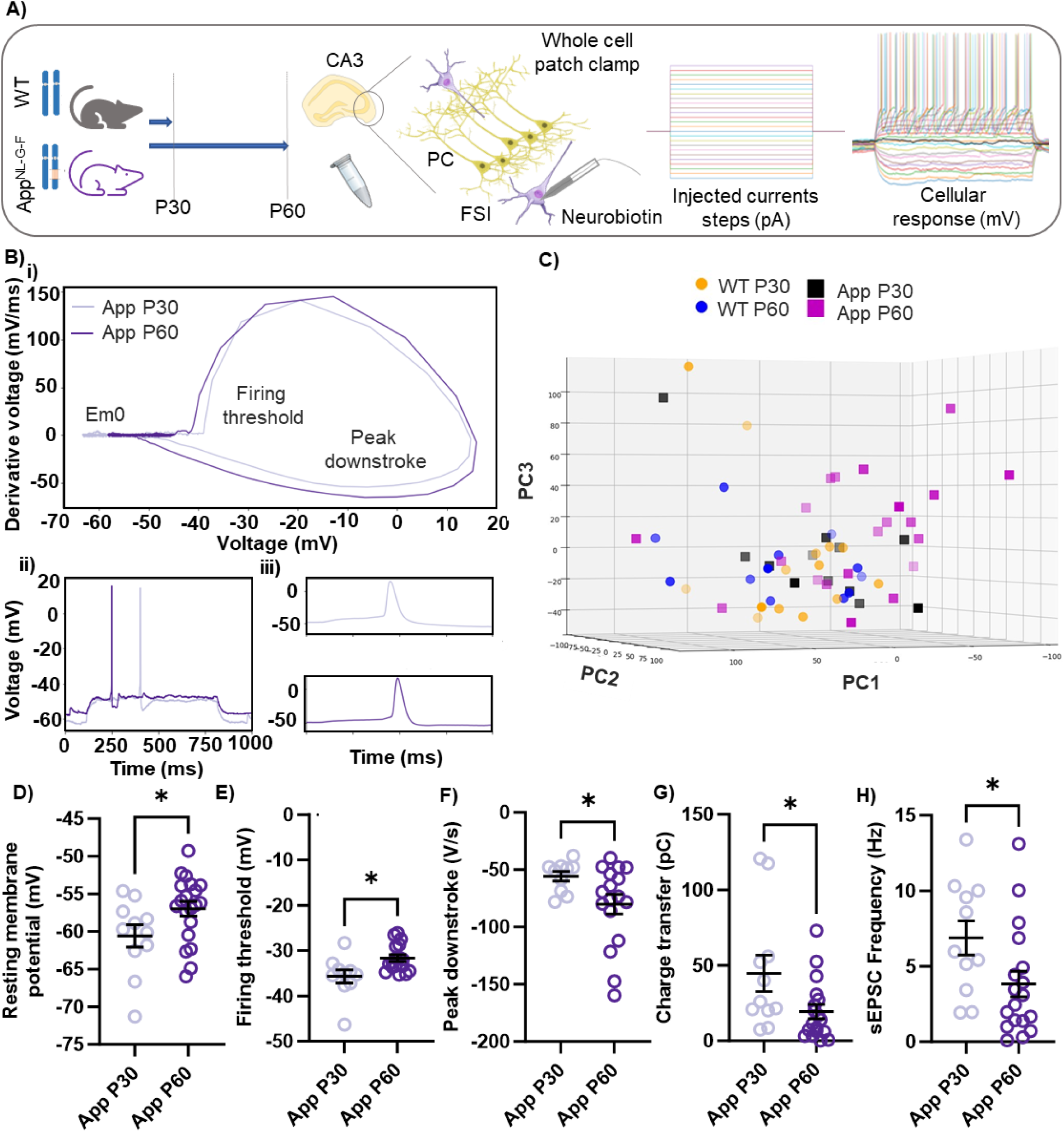
Fast-spiking interneurons intrinsic properties and postsynaptic input change during amyloid pathology progression. **A)** Graphic methods for whole cell patch clamp recordings from FSI. **B.i)** Representative phase-plot of the FSI’s action potentials from *App^NL-G-F^* mice at P30 and P60 showing different action potential shape and resting membrane potential (Em0) in the two groups. **B.ii)** Current-clamp recording traces of the representative FSI in B.i. **B.iii)** Cropped recording of the action potential of the FSI in Bi-ii. **C)** PCA of the intrinsic properties of WT P30 (n= 17), WT P60 (n= 11), *App^NL-G-F^* P30 (n=10), and *App^NL-G-F^* P60 (n=17) FSI. PC1 accounts for 37% of variation, PC2 for 34% and PC3 for 19%. *App^NL-G-F^* P60 FSI exhibit distinct properties that make them cluster separately from the other groups. **D-H)** Scatter plots of unpaired t-test of FSI’s intrinsic properties and post-synaptic input in the *App^NL-G-F^* group (P30 n=11 vs P60 n=17), showing **D)** resting membrane potential (p=0,044), **E)** firing threshold (p=0,011), **F)** peak downstroke (p=0,048), **G)** charge transfer (p=0,030), **H)** sEPSC frequency (p=0,049). Data are presented as mean ± SEM, and “n” indicates the number of neurons. *p < 0,05, **p < 0,01, ***p < 0,001.

**Table 1.** Intrinsic properties of FSI. Intrinsic properties of FSI in WT P30 (n= 17), WT P60 (n=11), *App^NL-G-F^* P30 (n=11) and *App^NL-G-F^* P60 (n=17). Data are presented as mean ± SEM, and “n” indicates the number of cells.

|  | Rheobase<br>(pA) | Firing<br>latency<br>(ms) | Firing<br>threshold<br>(mV) | AP<br>amplitude<br>(mV) | AP Half-width<br>(ms) | Peak<br>upstroke<br>(V/s) | Peak<br>downstroke<br>(V/s) |
| --- | --- | --- | --- | --- | --- | --- | --- |
| WT P30<br>(n=17) | 43,13 ±<br>4,892 | 93,96 ±<br>12,98 | -35,92 ±<br>1,725 | 58,25 ±<br>2,495 | 1,14 ±<br>0,066 | 148,7 ±<br>9,367 | -63,60 ±<br>4,827 |
| WT P60<br>(n=11) | 53,64 ±<br>7,600 | 53,64 ±<br>7,660 | -35,84 ±<br>1,485 | 56,50 ±<br>1,498 | 1,25 ±<br>0,087 | 140,1 ±<br>7,019 | -58,32 ±<br>6,397 |
| App P30<br>(n=11) | 35,45 ±<br>5,934 | 179,5 ±<br>25,490 | -35,67 ±<br>1,437 | 48,48 ±<br>3,337 | 1,16 ±<br>0,095 | 128,1 ±<br>9,510 | -54,12 ±<br>4,141 |
| App P60<br>(n=17) | 49,1 ±<br>6,832 | 155,9 ±<br>32,040 | -31,67 ±<br>0,733 | 47,41 ±<br>2,631 | 0,97 ±<br>0,079 | 130,9 ±<br>7,506 | -80,08 ±<br>8,594 |

**Table 1 – Intrinsic properties of FSI**
|  | Rise rate | Fall rate | Rise time<br>(ms) | Firing<br>rate (Hz) | Em 0<br>(mV) | Input<br>resistanc<br>e (GΩ) | Sag (mv) |
| --- | --- | --- | --- | --- | --- | --- | --- |
| WT P30<br>(n=17) | 97,48 ±<br>6,367 | -38,35 ±<br>3,260 | 0,6211 ±<br>0,025 | 6,846 ±<br>0,768 | -57,57 ±<br>1,323 | 0,401 ±<br>0,060 | 3,685 ±<br>0,547 |
| WT P60<br>(n=11) | 87,47 ±<br>5,324 | -31,70 ±<br>4,143 | 0,6467 ±<br>0,033 | 7,070 ±<br>1,109 | -59,57 ±<br>1,516 | 0,342 ±<br>0,033 | 3,977 ±<br>0,410 |
| App P30<br>(n=11) | 83,54 ±<br>6,382 | -34,02 ±<br>3,732 | 0,590 ±<br>0,024 | 10,860 ±<br>1,437 | -60,60 ±<br>1,481 | 0,388 ±<br>0,035 | 5,457 ±<br>0,930 |
| App P60<br>(n=17) | 88,41 ±<br>5,302 | -45,59 ±<br>4,555 | 0,594 ±<br>0,040 | 8,594 ±<br>1,368 | -56,99 ±<br>0,974 | 0,370 ±<br>0,032 | 4,378 ±<br>0,570 |

To evaluate differences and similarities within FSI across all groups, we performed a three-dimensional principal component analysis (PCA) on the intrinsic properties of FSI (Table 1). This analysis revealed that FSI in the *App^NL-G-F^* group at P60 had distinctive intrinsic properties since they clustered in a separate area of the plot compared to the *App^NL-G-F^* group at P30 and the WT groups at P30 and P60 (Fig 2.C; supp. video 1). This result indicates that FSI in the *App^NL-G-F^* mice at P30 were functionally comparable to those from the WT groups, while at P60 they diverged from the WT groups and the *App^NL-G-F^* P30 group, implying an alteration of FSI intrinsic properties during amyloidosis progression (*App^NL-G-F^* P30 vs P60; Fig 2.B). Indeed, FSI in the *App^NL-G-F^* group at P60 were depolarized (p=0,044; Fig 2.D), and they exhibited a higher firing threshold (p=0,011; Fig 2.E) and faster repolarization phase of the action potential (p=0,049; Fig 2.F) in comparison with FSI in the *App^NL-G-F^* group at P30. Additionally, FSI from P60 *App^NL-G-F^* mice received a decreased postsynaptic input, as shown by charge transfer (p=0,027, Fig 2.G) and spontaneous excitatory postsynaptic current (sEPSC) frequency (p=0,027, Fig 2.H). These findings suggest a change in the FSI membrane properties during the amyloid pathology progression. Importantly, the properties that differed between *App^NL-G-F^* P60 and P30 were preserved in the *App^NL-G-F^* P30 group as compared to age-matched WT mice (Supp fig 2), confirming that the functionality of FSI was not affected at this time point.

Collectively, such changes in FSI functionality might contribute to the impairment of the hippocampal network (Fig 1.B, C) and the FSI spike-gamma uncoupling [29] that we observed in the early stage of the amyloid pathology progression.

### 3.3 Bulk RNA sequencing reveals synaptic-related transcriptional changes during the early stage of amyloid pathology progression

Next, we performed bulk RNA sequencing to investigate the molecular events before and after the impairment in the hippocampal oscillatory activity and neuronal properties in the *App^NL-G-F^* mouse model (Fig 3.A).

**Fig 3.**
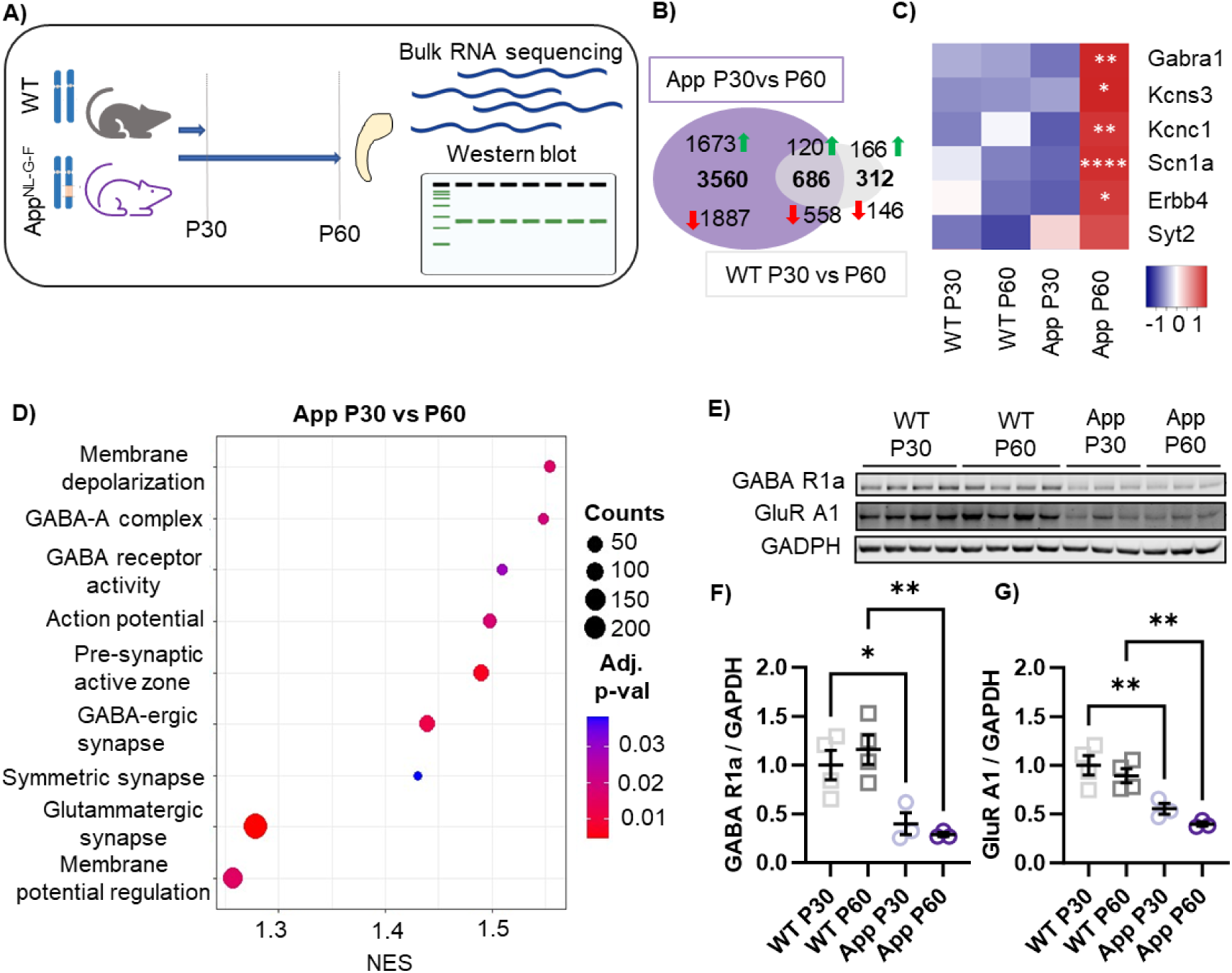
Bulk RNA sequencing reveals synaptic-related transcriptional changes during the early stage of amyloid pathology progression. **A)** Graphic methods for bulk RNA sequencing and western blot analysis from whole hippocampal tissue. **B)** Venn diagram showing unique and overlapping transcriptional changes between WT P30 vs P60 and *App^NL-G-F^* P30vs P60. The green arrows indicate upregulated genes and the red arrows indicate downregulated genes. **C)** Heat map showing the differences in the normalized count of selected genes relevant for FSI’s functionality and gamma oscillation onset in WT P30 (n=4), WT P60 (n=4), *App^NL-G-F^* P30 (n=4), *App^NL-G-F^* P60 (n=4). The significance level indicates the comparison of *App^NL-G-F^* P30 vs P60 of *Gabra1* (Log2FC=0,252, p= 0,002), *Kcns3* (Log2FC=0,216, p= 0,031), *Kcnc1* (Log2FC=0,282, p= 0,0002), *Scn1a* (Log2FC=0,393, p<0,0001), *Erbb4* (Log2FC=0,269, p= 0,016), *Syt2* (Log2FC=0,209, p= 0,118) obtained from the Wald test. **D)** Dot plot of the gene set enrichment analysis in *App^NL-G-F^* P30 vs P60, showing changes involving synaptic transmission and neuronal excitability regulation. Western blot membrane of GABA receptor 1 alpha, Glutamate receptor A1 and GADPH from VT P30 (n=4), WT P60 (n=4), *App^NL-G-F^* P30 (n=4), *App^NL-G-F^* P60 (n=4). **E)** Scatter plot of one-way ANOVA with Holm–Sidak’s multiple comparisons test of GABA R1a normalized expression showing a downregulation in *App^NL-G-F^* (WT vs *App^NL-G-F^* P30 p= 0,034; WT vs *App^NL-G-F^* P60 p= 0,005; *App^NL-G-F^* P30 vs P60, p=0,9466) **G)**. Scatter plot of one-way ANOVA with Holm–Sidak’s multiple comparisons test of GluR A1 protein level showing a decrease in *App^NL-G-F^* (WT vs *App^NL-G-F^* P30 p= 0,009; WT vs *App^NL-G-F^* P60 p=0,005; *App^NL-G-F^* P30 vs P60, p=0,561). Scatter plots data are presented as mean ± SEM, and “n” indicates the number of animals. *p < 0,05, **p < 0,01, ***p < 0,001, **** p < 0,0001.

First, we analyzed the dataset from WT mice at P30 vs P60, showing 998 differentially expressed genes (DEG) plausibly due to brain maturation (Supp fig 3.A). Then we compared age-matched WT vs *App^NL-G-F^* groups at P30 and P60 age to evaluate the amyloidogenic effect in the *App^NL-G-F^* knock-in model in comparison to WT mice. We found 510 DEG at P30 WT vs *App^NL-G-F^* (Supp fig 4.A) and 930 DEG at P60 WT vs *App^NL-G-F^* (Supp fig 4.B). Finally, to assess the effects of amyloid pathology progression we compared the *App^NL-G-F^* model at P30 vs P60 and we found 4246 DEG (Supp fig 3.B).

To differentiate the transcriptional changes triggered by amyloid pathology (App^NL-G-F^ P30 vs P60) from the ones due to normal brain maturation (WT P30 vs P60), we compared the DEGs from these two groups (Fig. 3.B). We found that 686 DEGs (120 upregulated and 588 downregulated; supp fig 5.A) were shared between the two comparisons, suggesting that it is genes that are indeed associated with brain maturation in both WT and *App^NL-G-F^* mice. Interestingly, 3560 DEG (1673 upregulated and 1887 downregulated, supp fig 5.A) changed exclusively during amyloid pathology progression (*App^NL-G-F^* P30 vs P60).

Next, to find the biological pathways involved in the transcriptional changes we performed an enrichment analysis (Supp. Table 2) at P30 (WT P30 vs *App^NL-G-F^* P30, supp fig 5.B), at P60 (WT P60 vs *App^NL-G-F^* P60, supp fig 5.C), during brain maturation (WT P30 vs P60, supp fig 5.D) and amyloid pathology progression (*App^NL-G-F^* P30 vs P60, fig 3.D). Our analysis reveals the activation of synaptic pathways in the *App^NL-G-F^* group compared to the WT at P30 (WT P30 vs *App^NL-G-F^* P30, Supp Fig 5.B). In contrast, at P60, the enrichment analysis shows fewer synaptic pathways when we compare the *App^NL-G-F^* and WT groups (WT P60 vs *App^NL-G-F^* P60, Supp Fig 5.C). Moreover, we found that the synaptic pathways were scarcely affected by maturation (WT P30 vs P60 Supp Fig 5.D). Interestingly, multiple pathways related to synaptic transmission, GABAergic and glutamatergic communication, ion channel activity, synaptic communication, and membrane potential and action potential regulation were activated during amyloid pathology progression (Fig 3.D). Candidate genes belonging to these pathways were selected based on their involvement in gamma oscillations and FSI functionality as potential causes of early functional impairment during amyloid pathology progression. *Kcnc1* [45–47]*, Kcns3* [48,49]*, Scn1a* [50,51]*, Erbb4* [52–55]*, GABRA1* [56,57] and *Syt2* [58–61] were upregulated in the *App^NL-G-F^* group at P60 (Fig 3.C; Supp table 1).

After, we aimed to investigate whether the transcriptional changes during amyloidosis pathology progression detected by RNA sequencing translated to protein level changes (*App^NL-G-F^* P30 vs P60). For this purpose, we quantified specific proteins corresponding to the altered genes in our transcriptomic assessment (ErbB4, KCNS3, Nav 1.1, Kv 3.1, Syt 2, GABA A b3, and GluR4) and some additional receptor subunits involved in the FSI and gamma oscillations activity: Glutamate receptor A1, GABA receptor A beta3 [31] and glutamate receptor 4 [62]. Our results showed that GABA receptor 1 alpha (Fig. 3.E, F) and glutamate receptor A1 (Fig. 3.E, G) were significantly downregulated already at P30 in the *App^NL-G-F^* mice when compared to WT mice, and the downregulation is also detected at P60. No other changes in protein levels were detected at these time points (Supp Fig 6).

### 3.4 Microglia-related transcriptional changes indicate loss of homeostatic and inflammatory microglia during the early stage of amyloid pathology progression

Given the crucial role of microglia in neuronal network regulation [21] and AD [7,63], our next step was to investigate transcriptional changes of microglia-related genes and pathways.

Pathway enrichment analysis (Supp. Table 2) showed that microglia-and inflammation-related processes, including microglia differentiation, acute inflammatory response, phagocytosis, synaptic pruning, and complement systems were suppressed in the *App^NL-G-F^* group at P60 compared to P30 (*App^NL-G-F^* P30 vs P60; Supp fig 7.A). Interestingly, similar pathways are activated in *App^NL-G-F^* mice at P30 compared to age-matched WT mice (WT vs *App^NL-G-F^* P30; Supp fig 7.B). Fewer microglia and inflammation-related changes occur with normal brain maturation in the WT group (WT P30 vs P60; Supp fig 7.C) and in the *App^NL-G-F^* group at P60 compared to age-matched WT mice (WT vs *App^NL-G-F^* P60; Supp fig 7.D). These transcriptional changes suggest a general activation of microglia and inflammation-related pathways in the *App^NL-G-F^* group at P30, followed by their suppression at P60.

Furthermore, we evaluated the transcriptional differences of microglia populations across the four groups using a two-dimensional PCA. We selected 2054 microglia genes, representative of homeostatic and inflammatory microglia [64–66] from the RNA sequencing dataset (Supp. Table 3). The PCA underlines the similarity of the microglia expression profiles in the WT group at P30 and P60. Conversely, microglia gene expression differs between P30 and P60 in the *App^NL-G-F^* mice, with samples from P30 and P60 time points clustering separately at the opposite edges of the PCA plot (Fig. 4.A). A similar sample distribution can be observed when genes representative of single microglia populations are plotted separately (Homeostatic and pruning microglia, 751 genes, Supp fig 8.A; activated microglia, 885 genes, Supp fig 8.B; interferon responding microglia, 418 genes, Supp fig 8.C). This result suggests that the microglia transcriptional profile changes to a higher extent during amyloid pathology progression than it would during normal maturation.

**Fig 4.**
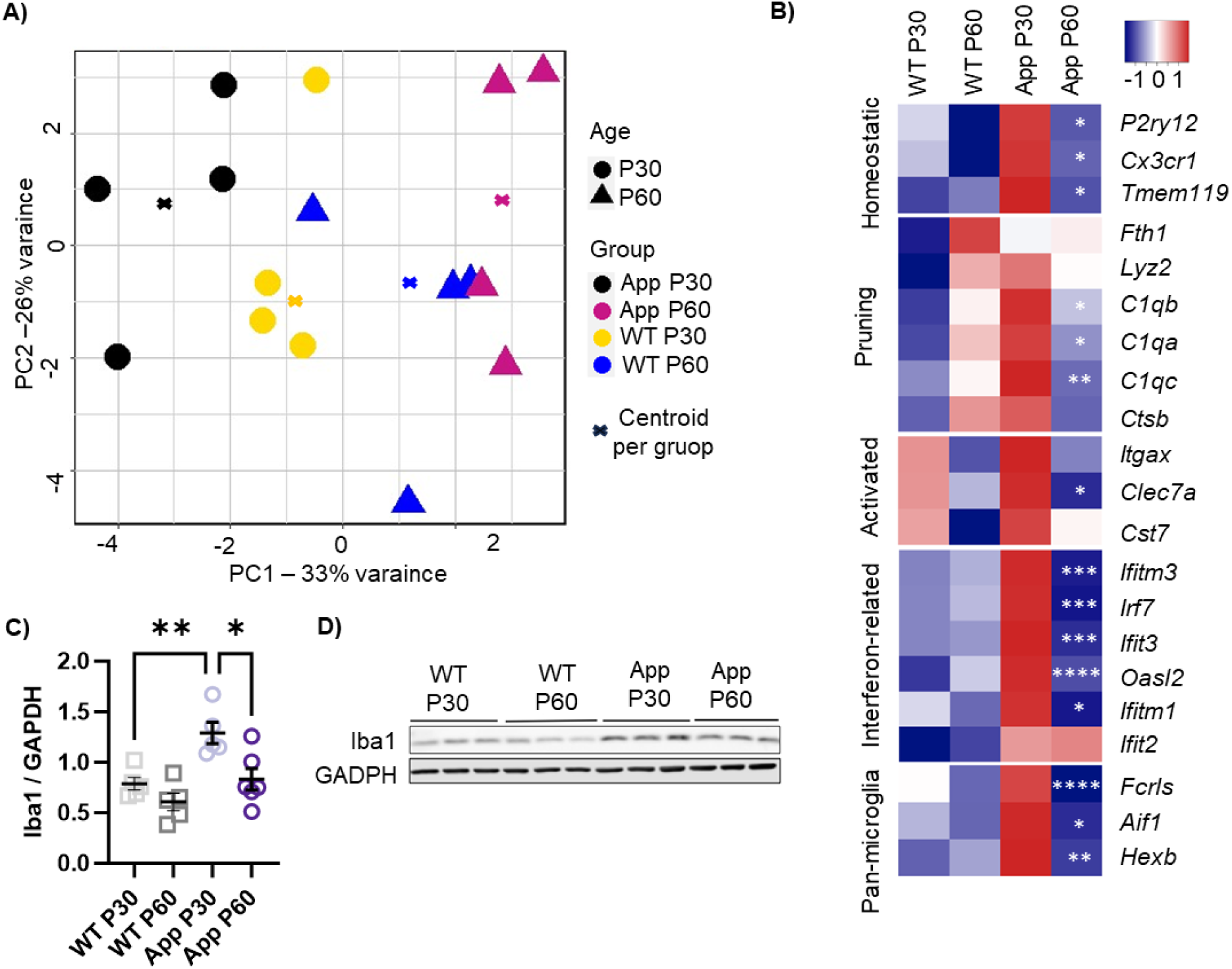
Microglia-related transcriptional changes indicate loss of homeostatic and inflammatory microglia during the early stage of amyloid pathology progression. **A)** PCA of 2054 microglia-related gene expression in WT P30 (n= 4), WT P60 (n= 4), *App^NL-G-F^* P30 (n=4), *App^NL-G-F^* P60 (n=4). Centroids represent the geometrical centre of each group. **B)** Heat maps showing the differences in the normalized count of microglial genes associated with different microglia phenotypes in WT P30 (n=4), WT P60 (n=4), *App^NL-G-F^* P30 (n=4), *App^NL-G-F^* P60 (n=4). The significance level indicates the comparison of *App^NL-G-F^* P30 vs P60 of *P2ry12* (Log2FC= -0,205, p= 0,031), *Cx3cr1* (Log2FC= -0,206, p= 0,026) and *Tmem119* (Log2FC= -0,213, p= 0,023), for homeostatic microglia; *Fth1* (Log2FC= -0,020, p= 0,872), *Lyz2* (Log2FC= -0,054, p= 0,657), *C1qb* (Log2FC= -0,196, p= 0,004), *C1qa* (Log2FC= -0,216, p= 0,031), *C1qc* (Log2FC= -0,270, p= 0,002), *Ctsb* (Log2FC= -0,082, p= 0,100) for homeostatic microglia engaged in synaptic pruning; *Itgax* (Log2FC= -0,097, p= 0,150), *Clec7a* (Log2FC= -0,329, p= 0,035), *Cst* (Log2FC= 0,158, p= 0,33) for activated microglia; *Iftitm3* (Log2FC= -0,593, p= 0,0001), *Irf7* (Log2FC= -0,745, p= 0,0005), *Ifit3* (Log2FC= -0,560, p= 0,0009), *Oasl2* (Log2FC= -0,649, p<0,00001), *Ifitm1* (Log2FC= -0,681, p= 0,011), *Ifit2* (Log2FC= 0,078, p= 0,956) for interferon related microglia; *Fcrls* (Log2FC= -0,448, p<0,00001), *Aif1* (Log2FC= -0,257, p= 0,018), *Hexb* (Log2FC= -0,234, p= 0,009) for pan-microglia. **C)** Scatter plot of one-way ANOVA with Holm–Sidak’s multiple comparisons test of Iba1 normalized protein level showing an increase in *App^NL-G-F^* at P30 (WT n=5 vs *App^NL-G-F^* n=5, p= 0,009) and a decrease in *App^NL-G-F^* at P60 (*App^NL-G-F^* P30 n=5 vs P60 n=6, p= 0,011). **D)** Representative western blot membrane of Iba1 and GADPH from WT P30 (n=3), P60 (n=3), and *App^NL-G-F^* P30 (n=3), P60 (n=3). Scatter plot data are presented as mean ± SEM, and “n” indicates the number of animals. *p < 0,05, **p < 0,01, ***p < 0,001, **** p < 0,0001.

To further characterize this event, we analyzed the expression of distinctive genes for different subpopulations of microglia previously reported by single-nucleus RNA sequencing characterization of microglia in the *App^NL-G-F^* mouse model [64]: *Tmem119, P2ry12,* and *Cx3cr1* for homeostatic microglia; *C1qa, C1qb, C1qc, Ctsb, Ctsd, Fth1* and *Lyz2* homeostatic microglia engaged in synaptic pruning; *Irf7, Ifitm3, Ifit3, Oasl2, Ifitm1* and *Ifit2* for interferon responding microglia; *Clec7a, Cst7* and *Itgax* for activated response microglia. A significant downregulation of genes related to microglia homeostatic and inflammatory states was detected in the *App^NL-G-F^* group at P60 compared to P30 (*App^NL-G-F^* P30 vs P60; Fig 4.B). Additionally, widely accepted pan-microglia markers such as *Aif1, Fcrls,* and *Hexb* [67] followed the same expression pattern, suggesting a suppression of microglia-related genes in *App^NL-G-F^* mice at P60 that could indicate a loss of microglia at this time point. To test this hypothesis, we measured the protein levels of Iba1, which is the protein encoded by *Aif1* and a commonly used pan-microglia marker, by western blot (Fig 4.C, D). The level of Iba1 increased in P30 *App^NL-G-F^* mice compared to age-matched WT mice (WT vs *App^NL-G-F^* P30, p=0,009; fig 4.C) and decreased between P30 and P60 in the *App^NL-G-F^* group (*App^NL-G-F^* P30 vs P60, p=0,013; fig 4.C).

### 3.5 Iba1-positive microglia density decreases during the early stage of amyloid pathology and correlates with plaque aggregation and FSI dysfunction

To verify that the microglia-related transcriptional and protein level variations reflected a change in the density of microglia in the hippocampus of *App^NL-G-F^* mice, we performed immunofluorescence staining (Fig 5.A-D) of hippocampal slices containing neurobiotin-marked FSI from our intracellular recordings (Fig 5.I, 2.A).

**Fig 5.**
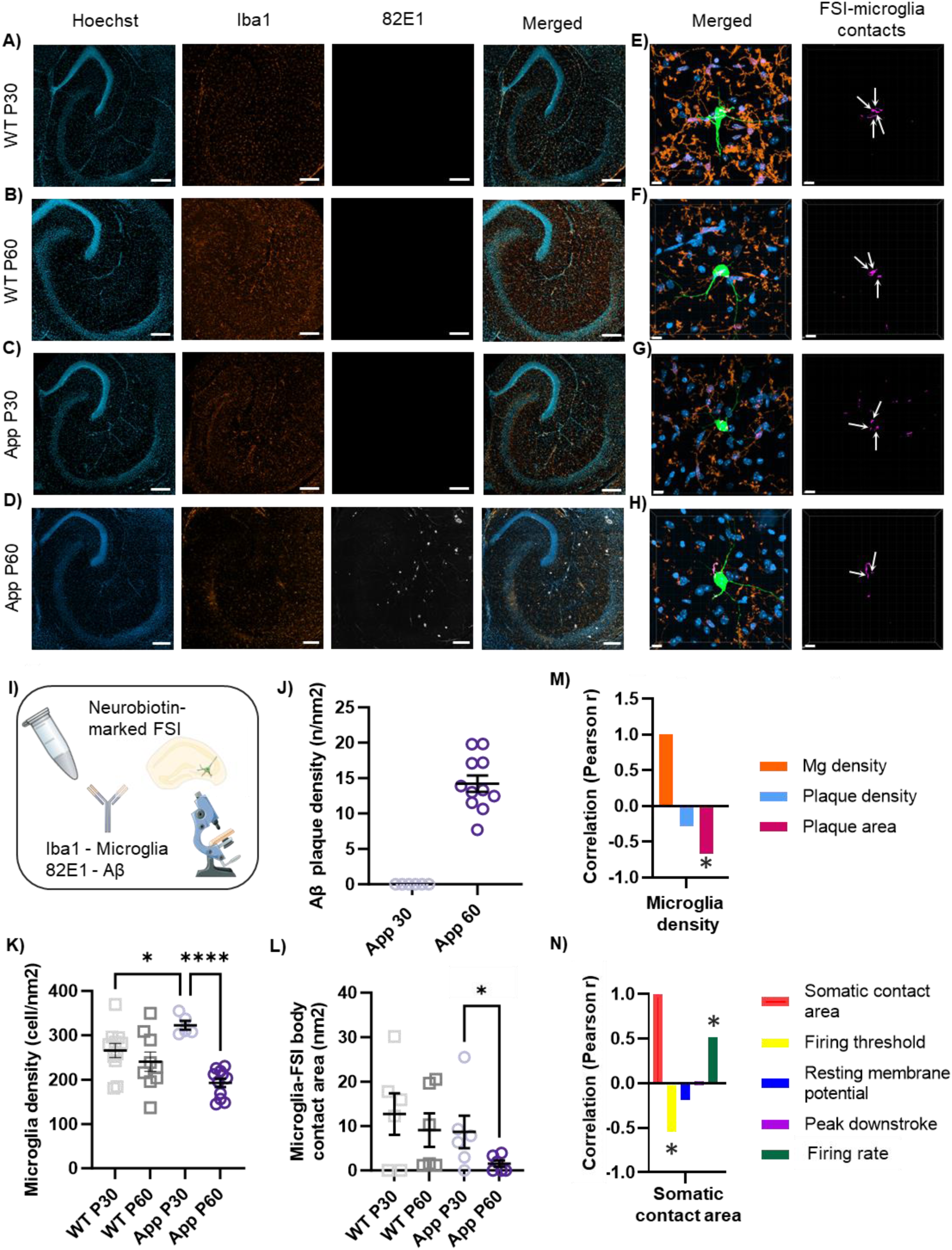
Iba1-positive microglia density decreases during the early stage of amyloid pathology and correlates with plaque aggregation and FSI dysfunction. **A-D)** Representative immunofluorescent staining of nuclei (Hoechst - blue), microglia (Iba1-red), Aβ (82E1-white) in the hippocampus of **A)** WT P30 (n= 11), **B)** WT P60 (n=9), **C)** *App^NL-G-F^* P30 (n=6) and **D)** *App^NL-G-F^* P30 (n=11). **E-H)** Imaris 3D reconstruction of neurobiotin-marked FSI (green), microglia (Iba1-red), nuclei (Hoechs-blue) and FSI-microglia contacts (pink) in **E)** WT P30 (n= 6), **F)** WT P60 (n=6), **G)** *App^NL-G-F^* P30 (n=6) and **H)** *App^NL-G-F^* P30 (n=6). **I)** Graphic method for immunofluorescent staining. **J)** Scatter-plot showing Aβ plaque density in the hippocampus of *App^NL-G-F^* P30 and *App^NL-G-F^* P60. **K)** Scatter-plot of one-way ANOVA with Holm–Sidak’s multiple comparisons test of microglia density, which increases in *App^NL-G-F^* P30 (WT P30 vs *App^NL-G-F^* P30 p= 0,036) and decreases in *App^NL-G-F^* P60 (WT P60 vs *App^NL-G-F^* P30 p= 0,065, *App^NL-G-F^* P30 vs *App^NL-G-F^* P60 p<0,0001). **L)** Scatter-plot of one-tail unpaired t-test of FSI-microglia somatic contact area in WT and *App^NL-G-F^* groups showing a decrease in *App^NL-G-F^* P60 (*App^NL-G-F^* P30 vs P60, p=0,0418). **M)** Bar-plot of Pearson correlation test of microglia density with Aβ plaque density and Aβ plaque area in the hippocampus of *App^NL-G-F^* P60. Microglia density inversely correlates with Aβ plaque area (correlation score r =-0,671; p=0,048). **N)** Bar-plot of Pearson correlation test of microglia-FSI somatic contact area and FSI firing threshold, resting membrane potential, peak downstroke, and firing frequency of FSI regardless of the group (n=19). Microglia-FSI somatic contact area inversely correlates with FSI firing threshold (correlation score r=-0,548; p=0,015) and directly correlates with FSI’s firing frequency (correlation score r=0,519; p=0,023). Scatter-plot data are presented as mean ± SEM, “n” indicates the number of slices in A-D, L, N and the number of neurons in E-G, M, O. Scale-bar represents 200μm in A-D and 10μm in E-H. *p < 0,05, **p < 0,01, ***p < 0,001, **** p < 0,0001.

Here we showed that microglia density, expressed as the number of Iba1-positive cells by area, increased at P30 in *App^NL-G-F^* mice compared to age-matched WT mice (WT vs *App^NL-G-F^* P30, p=0.036; fig 5.K). Importantly, we found a decrease in the hippocampus of *App^NL-G-F^* mice at P60 when compared to P30 (*App^NL-G-F^* P30 vs P60, p<0,0001; fig 5.K). These findings validate the dynamic pattern of microglia abundance and gene expression observed by the transcriptomic and protein analysis.

Besides microglia, we stained Aβ to evaluate the stage of plaque development during amyloid pathology progression in the hippocampus. As expected, the hippocampi of WT mice were free of amyloid plaques (Fig 5.A,B). In contrast, we found small, non-widespread amyloid plaques in the hippocampi of *App^NL-G-F^* mice at P60, but not at P30 (Fig 5.C-D, J). Interestingly, the area occupied by amyloid plaques inversely correlated with the density of Iba1-positive microglia in the hippocampus of P60 *App^NL-G-F^* mice (Fig 5.M), suggesting that the loss of microglia could play a role in plaque aggregation at this stage of the pathology. Additionally, to investigate whether the loss of Iba1-positive microglia in P60 *App^NL-G-F^* mice affected the interaction between microglia and FSI, we used Imaris to obtain a 3D reconstruction of neurobiotin-marked FSI, labeled during patch-clamp recordings, and microglia (Fig 5.E-H). We found that the somatic contact area between microglia and FSI is decreased at P60 when compared to P30 in *App^NL-G-F^* mice (*App^NL-G-F^* P30 vs P60, p=0,069; fig 5.L), while the number of contacts is similar between *App^NL-G-F^* P30 and WT P30 and P60 (Fig 5.L).

Since somatic microglia-neuron interaction has been reported to regulate neuronal activity and excitability [22,68], we explored a potential connection between the area of microglia-FSI somatic interphase and FSI excitability in all the groups. To this end, we tested the correlation between microglia-FSI contact area and the FSI intrinsic properties that were affected in P60 *App^NL-G-F^* mice (Fig 2.D-F), adding the firing rate as an additional parameter to evaluate excitability. Interestingly, the microglia-FSI somatic contact area positively correlated with FSI firing rate and negatively with their firing threshold, indicating that FSI with a smaller microglia-FSI somatic interphase area had a higher firing threshold (p=0,015) and fired at a lower rate (p=0,023, Fig 5.N).

Taken together, our results show a higher density of Iba1-positive microglia in the hippocampus of *App^NL-G-F^* mice compared to WT before plaque deposition and functional impairment, which decreases one month later, in correspondence with the beginning of amyloid plaque aggregation and network dysfunction. In addition, this loss of microglia in the *App^NL-G-F^* group at P60 is associated with changes in the firing activity of FSI, through a lack of microglia-FSI somatic contacts.

## 4 Discussion

In this study, we show that amyloid pathology in *App^NL-G-F^* mice progressively and detrimentally affects the hippocampal circuit responsible for gamma oscillations to a point of network failure, before widespread Aβ plaque load. Concurrently, the intrinsic properties of FSI are compromised during this early stage of the pathology. Interestingly, we found that this early functional impairment is associated with a loss of Iba1-positive microglia in the hippocampus and a decrease in their somatic contacts with FSI. We propose that this loss of microglia could be connected to the onset of Aβ plaque aggregation and network dysfunction in the hippocampus.

Increasing evidence has shown a crucial role of microglia in AD pathogenesis and progression [7,63]. However, the involvement of this cell type in AD pathology development and its role in Aβ plaque formation and clearance remains controversial [7,11]. One of the main difficulties to overcome when studying microglia is their extreme dynamism, the reason why different studies can lead to apparently contradictory results when trying to shed light on their role in AD pathophysiology. Studies have shown the protective function of microglia activation at pre-symptomatic stages of amyloidosis both in animal models [12,15] and clinical studies [13,17]. Indeed, in patients in a preclinical stage of AD, microglia activation correlated with a slower cognitive decline [13] and amyloid deposition [14]. Recently, Yang et al. reported that priming microglia before plaque aggregation promotes Aβ phagocytosis, reduces the size of Aβ plaques, and improves memory performance in 5xFAD mice [15].

On the other hand, the contribution of microglia to plaque formation and compaction has been hypothesized [69–71]. In particular, Spangenberg et al. reported that microglia depletion impairs Aβ plaque formation in 5xFAD mice [71]. The mechanism behind this process remains unclear. One hypothesis is that excessive accumulation of Aβ in microglia lysosomes leads to cell death, which might promote plaque aggregation by the release of Aβ agglomerates at the site of microglial death [72]. Additionally, strong evidence suggests that microglia reactivity promotes neurodegeneration by sustaining chronic inflammation at advanced AD stages [17,19]. In agreement with all these findings, a biphasic model has been proposed to explain microglia changes during AD progression. First, a protective peak of microglia activation arises in an early pre-symptomatic stage of the pathology, before Aβ plaque formation. Then a second detrimental activation phase promoting neuroinflammation takes place [14,63,73].

In this study, we found an increased density of Iba1-positive microglia in the hippocampus of *App^NL-G-F^* mice compared to WT (Fig 5.K) at a timepoint (P30) that precedes Aβ plaque aggregation (Fig 5.C, J) and functional impairment (Fig 1.B-C). An increased number of microglia cells is considered a sign of microglia activation during inflammatory response [74,75], in line with studies reporting an inflammatory response of microglia before plaque formation in animal models of amyloid pathology [18,76–78]. This suggests microglia activation in response to Aβ clearing, which keeps the brain free of Aβ plaques and maintains the hippocampal synaptic function.

However, as the pathology progresses in the *App^NL-G-F^* mice (P60), we observe a loss of Iba1-positive microglia in comparison to the initial increase (P30; Fig 5.K). This takes place concurrently with the onset of functional impairment (Fig 1.B) and Aβ plaque aggregation (Fig 5.D, J) in the hippocampus. Moreover, homeostatic and inflammatory microglia genes [64] are downregulated at P60 compared to P30 (Fig 4.B), suggesting that the reduction in the number affects in general the diverse microglia populations. Furthermore, in our samples the density of Iba1-positive microglia in the hippocampus negatively correlated with the hippocampal area occupied by Aβ plaques (Fig 5.M), suggesting that microglia loss might facilitate Aβ plaque aggregation at this time point.

While late microglia states in the *App^NL-G-F^* mouse model have been observed and characterized [64], this is the first time that an increased density of microglia is detected in a pre-plaque stage in this model. Although the causes and mechanisms behind the switch from protective to detrimental microglia remain to be elucidated [7], our findings of microglia loss between the two activation peaks could represent a turning point in this dynamic process.

Furthermore, we show that this decrease in Iba-1 positive microglia affects not only the total microglia number in the hippocampus but, importantly, its somatic contacts with FSI, which are also decreased (Fig 5.L). Microglia dynamically interact with neurons, constantly monitoring and regulating their activity [79]. Somatic junctions between microglia and neurons play a key role in this homeostatic communication [22,80]. Interestingly, in addition to decreased somatic contacts between microglia and FSI, we observed a downregulation of *P2y12r* (Fig 4.B), a purinergic receptor that is fundamental for microglia-neuron crosstalk [22], suggesting a loss of homeostatic surveillance of microglia on FSI. When somatic microglia-neuron interaction is compromised, changes in neuronal activity and excitability have been reported [22,68]. Consistently, our data show a change in FSI’s membrane and firing properties and a decrease in their postsynaptic input (Fig 2).

We found that microglia-FSI somatic contact area correlated positively with FSI firing rate and negatively with FSI firing threshold, which implies that FSI with smaller somatic contacts with microglia had a higher firing threshold and a lower firing rate (Fig 5.N). Our findings support the role of microglia surveillance in regulating FSI homeostasis, suggesting that a change in microglia-FSI somatic contacts can affect FSI excitability.

FSI are a subpopulation of inhibitory interneurons that, due to their specific firing properties [81–83], synchronize the hippocampal network in the gamma rhythm (20-80 Hz) [42,44,84]. Gamma oscillations are associated with higher cognitive functions [27], specifically hippocampal gamma oscillations are involved in memory formation and learning processes [26,28]. Alterations in gamma oscillation onset are related to cognitive deficits in multiple neurodegenerative diseases, including AD [85]. High concentrations of Aβ were reported to degrade hippocampal gamma oscillations in an acute amyloidogenic model [86]. Furthermore, we have proven that amyloid pathology leads to FSI desynchronization and gamma oscillation disruption, before widespread Aβ plaque deposition in the hippocampus of *App^NL-G-F^* mice [29]. We have now shown that resting membrane potential, firing threshold, action potential repolarization phase as well as postsynaptic current frequency and amplitude (Fig 2) of FSI change during amyloid pathology progression. These neuronal properties are crucial to regulate the intrinsic excitability of FSI [87], hence their modification can explain the FSI loss of rhythmicity and the consequent network failure that we observed in the hippocampus of *App^NL-G-F^* mice at P75 in our previous study [29]. Interestingly, restoration of FSI firing properties and rhythmicity leads to an improvement of gamma oscillation synchrony and cognitive performance [29,31,62].

Our transcriptomic analysis revealed that multiple pathways and genes related to synaptic transmission and neuronal excitability regulation are activated in correspondence with the hippocampal network dysfunction in the *App^NL-G-F^* model (Fig 3.C, D). Additionally, the protein expression of GABA receptor 1 alpha (Fig 3. F) and glutamate receptor A1 protein levels were decreased in the App^NL-G-F^ group at P30 and 60 (Fig 3.G). These receptor subunits belong to the main inhibitory and excitatory receptors expressed in the hippocampus, namely GABA A [56] and AMPA [88]. Given the importance of AMPA transmission for synaptic plasticity, this finding could explain the lack of increased IEG expression in App^NL-G-F^ mice upon KA-induced neuronal activation (Fig. 1.C, D) [89].

Here we propose that microglia play a protective role at earlier stages of amyloidosis preventing plaque aggregation and supporting FSI physiological functionality, which results in a preserved network synchronization in the gamma rhythm. Conversely, microglia loss in the early stage of amyloid pathology progression might facilitate plaque aggregation, cause functional impairment in FSI, and, consequently, prevent the synchronization of the hippocampal network in the gamma rhythm.

## 5 Conclusions

In conclusion, here we identified for the first time a loss of microglia during the early stage of amyloid pathology, before the manifestation of widespread Aβ plaque load in the hippocampus. Our findings suggest that microglia loss might represent a turning point in AD pathology progression, contributing to Aβ plaque formation, FSI dysfunction, and circuit failure in the hippocampus.

## Declarations

### Ethical approval

Experiments were performed in accordance with the ethical permit granted by Norra Stockholm’s Djurförsöksetiska Nämnd (dnr N45/13) to AF and 12570-2021 to PN.

### Consent for publication

Not applicable

### Availability of data and materials

Raw data is available from the corresponding authors upon request.

### Authors contributions

LEAG and AF designed the experiments. GP, LEAG, JG, CAR, SL, PRR, EP, GVC, MLL, FE, SM, PN, AFr, and KB performed experiments and analyses. GP created the Python scripts for analysis. GP and LEAG wrote the paper with inputs from all co-authors.

## Acknowledgments

We thank Simon Sundström for the guidance and help with the Python scripts and Johanna Wanngren for the help with mouse breeding. We thank Takaomi Saido and Takashi Saito at RIKEN Center for Brain Science, Japan for providing App knock-in mice.

## References

1. Aisen PS, Cummings J, Jack CR, Morris JC, Sperling R, Frölich L, et al. On the path to 2025: Understanding the Alzheimer’s disease continuum. Alzheimers Res Ther. BioMed Central Ltd.; 2017.

2. Golde TE. Disease-Modifying Therapies for Alzheimer’s Disease: More Questions than Answers. Neurotherapeutics. Springer Science and Business Media Deutschland GmbH; 2022. p. 209–27.

3. Mehta D, Jackson R, Paul G, Shi J, Sabbagh M. Why do trials for Alzheimer’s disease drugs keep failing? A discontinued drug perspective for 2010-2015. Expert Opin Investig Drugs. 2017;26:735–9.

4. Braak H, Braak E. Neuropathological stageing of Alzheimer-related changes. Acta Neuropathol. 1991;82:239–59.

5. Hardy J, Selkoe DJ. The Amyloid Hypothesis of Alzheimer’s Disease: Progress and Problems on the Road to Therapeutics. Science (1979) [Internet]. 2022;297:353–6. Available from: https://www.science.org

6. Herrup K. The case for rejecting the amyloid cascade hypothesis. Nat Neurosci. 2015;18:794–9.

7. Leng F, Edison P. Neuroinflammation and microglial activation in Alzheimer disease: where do we go from here? Nat Rev Neurol. Nature Research; 2021. p. 157–72.

8. Passamonti L, Tsvetanov KA, Jones PS, Bevan-Jones WR, Arnold R, Borchert RJ, et al. Neuroinflammation and functional connectivity in Alzheimer’s disease: Interactive influences on cognitive performance. Journal of Neuroscience. 2019;39:7218–26.

9. Holmes C. Comment Common infections and increased risk of developing dementia : compelling evidence for intervention studies. Lancet Healthy Longev. 2021;2:e391–2.

10. Bohn B, Lutsey PL, Misialek JR, Walker KA, Brown CH, Hughes TM, et al. Incidence of Dementia Following Hospitalization With Infection Among Adults in the Atherosclerosis Risk in Communities (ARIC) Study Cohort. JAMA Netw Open. 2023;6:e2250126.

11. Hamelin L, Lagarde J, Dorothée G, Potier MC, Corlier F, Kuhnast B, et al. Distinct dynamic profiles of microglial activation are associated with progression of Alzheimer’s disease. Brain. 2018;141:1855–70.

12. Feng W, Zhang Y, Wang Z, Xu H, Wu T, Marshall C, et al. Microglia prevent beta-Amyloid plaque formation in the early stage of an Alzheimer’s disease mouse model with suppression of glymphatic clearance. Alzheimers Res Ther. 2020;12:1–15.

13. Hamelin L, Lagarde J, Dorothée G, Leroy C, Labit M, Comley RA, et al. Early and protective microglial activation in Alzheimer’s disease: A prospective study using 18F-DPA-714 PET imaging. Brain. 2016;139:1252–64.

14. Fan Z, Brooks DJ, Okello A, Edison P. An early and late peak in microglial activation in Alzheimer’s disease trajectory. Brain. 2017;140:792–803.

15. Yang Y, García-Cruzado M, Zeng H, Camprubí-Ferrer L, Bahatyrevich-Kharitonik B, Bachiller S, et al. LPS priming before plaque deposition impedes microglial activation and restrains Aβ pathology in the 5xFAD mouse model of Alzheimer’s disease. Brain Behav Immun. 2023;113:228–47.

16. Femminella GD, Dani M, Wood M, Fan Z, Calsolaro V, Atkinson R, et al. Microglial activation in early Alzheimer trajectory is associated with higher gray matter volume. Neurology. 2019;92:E1331–43.

17. Fan Z, Aman Y, Ahmed I, Chetelat G, Landeau B, Ray Chaudhuri K, et al. Influence of microglial activation on neuronal function in Alzheimer’s and Parkinson’s disease dementia. Alzheimer’s and Dementia. 2015;11:608–621.e7.

18. Heneka MT, Sastre M, Dumitrescu-Ozimek L, Dewachter I, Walter J, Klockgether T, et al. Focal glial activation coincides with increased BACE1 activation and precedes amyloid plaque deposition in APP[V717I] transgenic mice. J Neuroinflammation. 2005;2:1–12.

19. Mass E, Jacome-Galarza CE, Blank T, Lazarov T, Durham BH, Ozkaya N, et al. A somatic mutation in erythro-myeloid progenitors causes neurodegenerative disease. Nature. 2017;549:389–93.

20. Karson MA, Tang AH, Milner TA, Alger BE. Synaptic cross talk between perisomatic-targeting interneuron classes expressing cholecystokinin and parvalbumin in hippocampus. Journal of Neuroscience. 2009;29:4140–54.

21. Guan A, Wang S, Huang A, Qiu C, Li Y, Li X, et al. The role of gamma oscillations in central nervous system diseases: Mechanism and treatment. Front Cell Neurosci. 2022;16:962967.

22. Cserép C, Pósfai B, Lénárt N, Fekete R, László ZI, Lele Z, et al. Microglia monitor and protect neuronal function through specialized somatic purinergic junctions. Science (1979) [Internet]. 2020;367:528–37. Available from: https://www.science.org

23. Cserép C, Pósfai B, Dénes Á. Shaping Neuronal Fate: Functional Heterogeneity of Direct Microglia-Neuron Interactions. Neuron. Cell Press; 2021. p. 222–40.

24. De Haan W, Van der Flier WM, Koene T, Smits LL, Scheltens P, Stam CJ. Disrupted modular brain dynamics reflect cognitive dysfunction in Alzheimer’s disease. Neuroimage. 2012;59:3085–93.

25. Smailovic U, Koenig T, Kåreholt I, Andersson T, Gregoric M, Winblad B, et al. Neurobiology of Aging Quantitative EEG power and synchronization correlate with Alzheimer’s disease CSF biomarkers. Neurobiol Aging. 2018;63:88–95.

26. Colgin LL, Moser EI. Gamma oscillations in the hippocampus. Physiology. 2010;25:319–29.

27. Jensen O, Kaiser J, Lachaux JP. Human gamma-frequency oscillations associated with attention and memory. Trends Neurosci. 2007;30:317–24.

28. Düzel E, Penny WD, Burgess N. Brain oscillations and memory. Curr Opin Neurobiol. 2010;20:143–9.

29. Arroyo-García LE, Isla AG, Andrade-Talavera Y, Balleza-Tapia Hugo, Raúl Loera-Valencia, Laura Alvarez-Jimenez, et al. Impaired spike-gamma coupling of area CA3 fast-spiking interneurons as the earliest functional impairment in the App NL-G-F mouse model of Alzheimer’s disease. Mol Psychiatry [Internet]. 2021;1–11. Available from: 10.1038/s41380-021-01257-0

30. Arroyo-García LE, Bachiller S, Ruiz R, Boza-Serrano A, Rodríguez-Moreno A, Deierborg T, et al. Targeting galectin-3 to counteract spike-phase uncoupling of fast-spiking interneurons to gamma oscillations in Alzheimer’s disease. Transl Neurodegener. 2023;12:6.

31. Emre C, Arroyo-García LE, Do K V., Jun B, Ohshima M, Alcalde SG, et al. Intranasal delivery of pro-resolving lipid mediators rescues memory and gamma oscillation impairment in App NL-G-F/NL-G-F mice. Commun Biol. 2022;5.

32. Saito T, Matsuba Y, Mihira N, Takano J, Nilsson P, Itohara S, et al. Single App knock-in mouse models of Alzheimer’s disease. Nat Neurosci. 2014;17:661–3.

33. Arroyo-García LE, Bachiller S, Ruiz R, Boza-Serrano A, Rodríguez-Moreno A, Deierborg T, et al. Targeting galectin-3 to counteract spike-phase uncoupling of fast-spiking interneurons to gamma oscillations in Alzheimer’s disease. Transl Neurodegener. 2023;12.

34. Balleza-Tapia H, Arroyo-García LE, Isla AG, Loera-Valencia R, Fisahn A. Functionally-distinct pyramidal cell subpopulations during gamma oscillations in mouse hippocampal area CA3. Prog Neurobiol. 2022;210.

35. Wu T, Hu E, Xu S, Chen M, Guo P, Dai Z, et al. clusterPro fi ler 4 . 0 : A universal enrichment tool for interpreting omics data clusterPro fi ler 4 . 0 : A universal enrichment tool for interpreting omics data. The Innovation. 2021;2:100141.

36. Among T, Clusters G, Yu G. clusterProfiler : an R Package for Comparing Biological. 2012;16:284–7.

37. Minatohara K, Akiyoshi M, Okuno H. Role of immediate-early genes in synaptic plasticity and neuronal ensembles underlying the memory trace. Front Mol Neurosci. 2016;8.

38. Yap EL, Greenberg ME. Activity-Regulated Transcription: Bridging the Gap between Neural Activity and Behavior. Neuron. Cell Press; 2018. p. 330–48.

39. Kim S, Kim H, Um JW. Synapse development organized by neuronal activity-regulated immediate-early genes. Exp Mol Med. Nature Publishing Group; 2018.

40. Gallo FT, Katche C, Morici JF, Medina JH, Weisstaub N V. Immediate early genes, memory and psychiatric disorders: Focus on c-Fos, Egr1 and Arc. Front Behav Neurosci. 2018;12.

41. Dickey CA, Gordon MN, Mason JE, Wilson NJ, Diamond DM, Guzowskià JF, et al. Amyloid suppresses induction of genes critical for memory consolidation in APP + PS1 transgenic mice. J Neurochem [Internet]. 2004;88:434–42. Available from: https://onlinelibrary.wiley.com/terms-and-conditions

42. Cardin JA, Carlén M, Meletis K, Knoblich U, Zhang F, Deisseroth K, et al. Driving fast-spiking cells induces gamma rhythm and controls sensory responses. Nature. 2009;459:663–7.

43. Li Y, Zhu K, Li N, Wang X, Xiao X, Li L, et al. Reversible GABAergic dysfunction involved in hippocampal hyperactivity predicts early-stage Alzheimer disease in a mouse model. Alzheimers Res Ther. 2021;13.

44. Hu H, Gan J, Jonas P. Fast-spiking, parvalbumin + GABAergic interneurons: From cellular design to microcircuit function. Science (1979) [Internet]. 2014;345:1255263. Available from: https://www.science.org

45. Labro AJ, Priest MF, Lacroix JJ, Snyders DJ, Bezanilla F. Kv 3.1 uses a timely resurgent K + current to secure action potential repolarization. Nat Commun. 2015;6.

46. Kaczmarek LK, Zhang Y. Kv3 channels: Enablers of rapid firing, neurotransmitter release, and neuronal endurance. Physiol Rev. 2017;97:1431–68.

47. Andrade-Talavera Y, Arroyo-García LE, Chen G, Johansson J, Fisahn A. Modulation of Kv3.1/Kv3.2 promotes gamma oscillations by rescuing Aβ-induced desynchronization of fast-spiking interneuron firing in an AD mouse model in vitro. Journal of Physiology. 2020;598:3711–25.

48. Georgiev D, González-Burgos G, Kikuchi M, Minabe Y, Lewis DA, Hashimoto T. Selective expression of KCNS3 potassium channel α-subunit in parvalbumin-containing GABA neurons in the human prefrontal cortex. PLoS One. 2012;7.

49. Miyamae T, Hashimoto T, Abraham M, Kawabata R, Koshikizawa S, Bian Y, et al. Kcns3 deficiency disrupts Parvalbumin neuron physiology in mouse prefrontal cortex: Implications for the pathophysiology of schizophrenia. Neurobiol Dis. 2021;155.

50. Martin MS, Dutt K, Papale LA, Dubé CM, Dutton SB, De Haan G, et al. Altered function of the SCN1A voltage-gated sodium channel leads to γ-aminobutyric acid-ergic (GABAergic) interneuron abnormalities. Journal of Biological Chemistry. 2010;285:9823–34.

51. Holmes GL, Bender AC, Wu EX, Scott RC, Lenck-Santini PP, Morse RP. Maturation of EEG oscillations in children with sodium channel mutations. Brain Dev. 2012;34:469–77.

52. Vullhorst D, Neddens J, Karavanova I, Tricoire L, Petralia RS, McBain CJ, et al. Selective expression of ErbB4 in interneurons, but not pyramidal cells, of the rodent hippocampus. Journal of Neuroscience. 2009;29:12255–64.

53. Robinson HL, Tan Z, Santiago-Marrero I, Arzola EP, Dong TV, Xiong WC, et al. Neuregulin 1 and ErbB4 Kinase Actively Regulate Sharp Wave Ripples in the Hippocampus. Journal of Neuroscience. 2022;42:390–404.

54. Tan Z, Robinson HL, Yin DM, Liu Y, Liu F, Wang H, et al. Dynamic ErbB4 Activity in Hippocampal-Prefrontal Synchrony and Top-Down Attention in Rodents. Neuron. 2018;98:380–393.e4.

55. Zhang H, Zhang L, Zhou D, He X, Wang D, Pan H, et al. Ablating ErbB4 in PV neurons attenuates synaptic and cognitive deficits in an animal model of Alzheimer’s disease. Neurobiol Dis. 2017;106:171–80.

56. Olsen RW, Sieghart W. GABAA receptors: Subtypes provide diversity of function and pharmacology. Neuropharmacology. 2009. p. 141–8.

57. Kujala J, Jung J, Bouvard S, Lecaignard F, Lothe A, Bouet R, et al. Gamma oscillations in V1 are correlated with GABAA receptor density: A multi-modal MEG and Flumazenil-PET study. Sci Rep. 2015;5.

58. Chen C, Arai I, Satterfield R, Young SM, Jonas P. Synaptotagmin 2 Is the Fast Ca2+ Sensor at a Central Inhibitory Synapse. Cell Rep. 2017;18:723–36.

59. Pang ZP, Melicoff E, Padgett D, Liu Y, Teich AF, Dickey BF, et al. Synaptotagmin-2 is essential for survival and contributes to Ca 2+ triggering of neurotransmitter release in central and neuromuscular synapses. Journal of Neuroscience. 2006;26:13493–504.

60. Sommeijer JP, Levelt CN. Synaptotagmin-2 is a reliable marker for parvalbumin positive inhibitory boutons in the mouse visual cortex. PLoS One. 2012;7.

61. Maximov A. Synaptotagmins. In: Squire R., editor. Encyclopedia of Neuroscience,. Academic Pres; 2009. p. 819–21.

62. Verret L, Mann EO, Hang GB, Barth AMI, Cobos I, Ho K, et al. Inhibitory Interneuron Deficit Links Altered Network Activity and Cognitive Dysfunction in Alzheimer Model. Cell. 2012;149:708–21.

63. Sarlus H, Heneka MT, Sarlus H, Heneka MT. Microglia in Alzheimer’s disease Microglia in Alzheimer’s disease. J Clin Invest. 2017;127:3240–9.

64. Frigerio CS, Wolfs L, Fattorelli N, Perry VH, Fiers M, Strooper B De, et al. The Major Risk Factors for Alzheimer’s Disease : Age, Sex, and Genes Modulate the Microglia Response to A b Plaques Resource The Major Risk Factors for Alzheimer’s Disease : Age, Sex, and Genes Modulate the Microglia Response to A b Plaques. 2019;1293–306.

65. Hammond TR, Dufort C, Dissing-Olesen L, Giera S, Young A, Wysoker A, et al. Single-Cell RNA Sequencing of Microglia throughout the Mouse Lifespan and in the Injured Brain Reveals Complex Cell-State Changes. Immunity. 2019;50:253–271.e6.

66. Sun N, Victor MB, Park YP, Xiong X, Scannail AN, Leary N, et al. Human microglial state dynamics in Alzheimer’s disease progression. Cell. 2023;186:4386–4403.e29.

67. Jurga AM, Paleczna M, Kuter KZ. Overview of General and Discriminating Markers of Differential Microglia Phenotypes. Front Cell Neurosci. 2020;14:1–18.

68. Szalay G, Martinecz B, Lénárt N, Környei Z, Orsolits B, Judák L, et al. Microglia protect against brain injury and their selective elimination dysregulates neuronal network activity after stroke. Nat Commun. 2016;7.

69. Huang Y, Happonen KE, Burrola PG, O’Connor C, Hah N, Huang L, et al. Microglia use TAM receptors to detect and engulf amyloid β plaques. Nat Immunol. 2021;22:586–94.

70. Casali BT, MacPherson KP, Reed-Geaghan EG, Landreth GE. Microglia depletion rapidly and reversibly alters amyloid pathology by modification of plaque compaction and morphologies. Neurobiol Dis. 2020;142:104956.

71. Spangenberg E, Severson PL, Hohsfield LA, Crapser J, Zhang J, Burton EA, et al. Sustained microglial depletion with CSF1R inhibitor impairs parenchymal plaque development in an Alzheimer’s disease model. Nat Commun. 2019;10:3758.

72. Baik SH, Kang S, Son SM, Mook-Jung I. Microglia contributes to plaque growth by cell death due to uptake of amyloid β in the brain of Alzheimer’s disease mouse model. Glia. 2016;64:2274–90.

73. Onuska KM. The Dual Role of Microglia in the Progression of Alzheimer’s Disease. J Neurosci. 2020;40:1608–10.

74. Mander PK, Jekabsone A, Brown GC. Microglia Proliferation Is Regulated by Hydrogen Peroxide from NADPH Oxidase. The Journal of Immunology. 2006;176:1046–52.

75. Sun J, Zheng JH, Zhao M, Lee S, Goldstein H. Increased In Vivo Activation of Microglia and Astrocytes in the Brains of Mice Transgenic for an Infectious R5 Human Immunodeficiency Virus Type 1 Provirus and for CD4-Specific Expression of Human Cyclin T1 in Response to Stimulation by Lipopolysaccharides. J Virol. 2008;82:5562– 72.

76. Ferretti MT, Bruno MA, Ducatenzeiler A, Klein WL, Cuello AC. Intracellular Aβ-oligomers and early inflammation in a model of Alzheimer’s disease. Neurobiol Aging. 2012;33:1329–42.

77. Wright AL, Zinn R, Hohensinn B, Konen LM, Beynon SB, Tan RP, et al. Neuroinflammation and Neuronal Loss Precede Aβ Plaque Deposition in the hAPP-J20 Mouse Model of Alzheimer’s Disease. PLoS One. 2013;8.

78. Spangenberg EE, Green KN. Inflammation in Alzheimer’s disease: Lessons learned from microglia-depletion models. Brain Behav Immun. 2017;61:1–11.

79. Badimon A, Strasburger HJ, Ayata P, Chen X, Nair A, Ikegami A, et al. Negative feedback control of neuronal activity by microglia. Nature [Internet]. 2020;586:417–23. Available from: 10.1038/s41586-020-2777-8

80. Pósfai B, Cserép C, Orsolits B, Dénes Á. New Insights into Microglia – Neuron Interactions : A Neuron’s Perspective. Neuroscience. 2019;405:103–17.

81. McBain CJ. Interneurons unbound. Nat Rev Neurosci. 2001;2:11–23.

82. Pelkey KA, Chittajallu R, Craig MT, Tricoire L, Wester JC, McBain CJ. Hippocampal gabaergic inhibitory interneurons. Physiol Rev. 2017;97:1619–747.

83. Freund TF, Buzsáki G. Interneurons of the Hippocampus. Hippocampus. 1996;6:347–470.

84. Tremblay R, Lee S, Rudy B. Review GABAergic Interneurons in the Neocortex : From Cellular Properties to Circuits. Neuron. 2016;91:260–92.

85. Mably AJ, Colgin LL. Gamma oscillations in cognitive disorders. Curr Opin Neurobiol. 2018;52:182–7.

86. Kurudenkandy FR, Zilberter M, Biverstål H, Presto J, Honcharenko D, Strömberg R, et al. Amyloid-β-induced action potential desynchronization and degradation of hippocampal gamma oscillations is prevented by interference with peptide conformation change and aggregation. Journal of Neuroscience. 2014;34:11416–25.

87. Klemz A, Wildner F, Tütüncü E, Gerevich Z. Regulation of Hippocampal Gamma Oscillations by Modulation of Intrinsic Neuronal Excitability. Front Neural Circuits. 2022;15.

88. Traynelis SF, Wollmuth LP, McBain CJ, Menniti FS, Vance KM, Ogden KK, et al. Glutamate receptor ion channels: Structure, regulation, and function. Pharmacol Rev. 2010. p. 405–96.

89. Rao VR, Pintchovski SA, Chin J, Peebles CL, Mitra S, Finkbeiner S. AMPA receptors regulate transcription of the plasticity-related immediate-early gene Arc. Nat Neurosci. 2006;9:887–95.

