## Supplementary information for "Association of microglia loss with hippocampal network impairments as a turning point in the amyloid pathology progression"

### Supplementary data

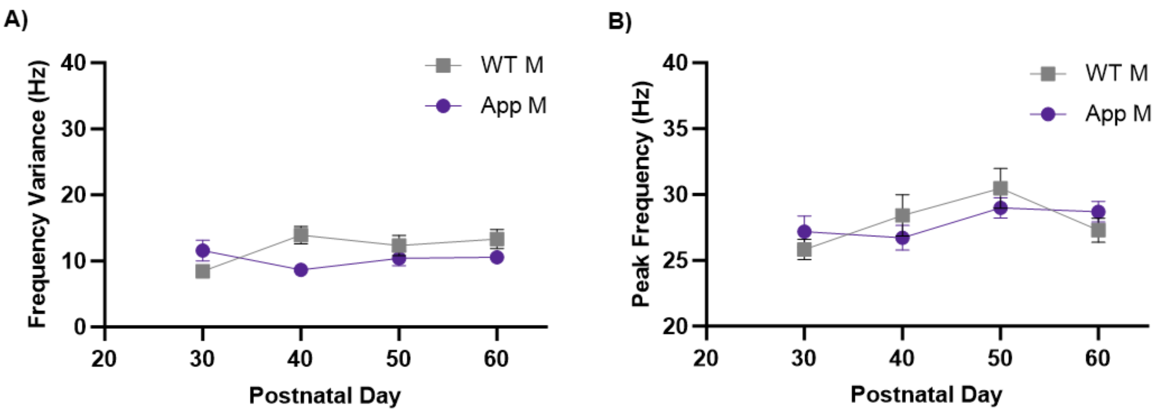


**Supplementary figure 1 – Frequency variance and peak frequency are stable during amyloid pathology and remain comparable with age-matched WT.**

**A-B)** Line plot of gamma **A)** frequency variance and **B)** peak frequency in WT and App$^{NL-G-F}$ mice at different ages. The graph shows a two-way ANOVA with Holm–Sidak's multiple comparisons test of **A)** frequency variance at P30 (WT n= 6 vs App$^{NL-G-F}$ n=6, p=0,831), P40 (WT n= 6 vs App$^{NL-G-F}$ n=6, p=0,1614), P50 (WT n= 6 vs App$^{NL-G-F}$ n=9, p=0,981), and P60 (WT n= 7 vs App$^{NL-G-F}$ n=11, p=0,802) and **B)** peak frequency at P30 (WT n= 6 vs App$^{NL-G-F}$ n=6, p=0,994), P40 (WT n= 6 vs App$^{NL-G-F}$ n=6, p=0,993), P50 (WT n= 6 vs App$^{NL-G-F}$ n=9, p=0,993), and P60 (WT n= 7 vs App$^{NL-G-F}$ n=11, p=0,993). Intra-group differences were evaluated too: for the **A)** frequency variance of WT group P30 vs P40 (p= 0,125), P30 vs P50 (p= 0,630), P30 vs P60 (p=0,202) and App$^{NL-G-F}$ group P30 vs P40 (p=0881), P30 vs P50 (p= 0,989), P30 vs P60 (p= 0,989). For the **B)** Peak frequency of WT group P30 vs P40 (p= 0,928), P30 vs P50 (p= 0,141), P30 vs P60 (p=0,993) and App$^{NL-G-F}$ group P30 vs P40 (p=0,992), P30 vs P50 (p= 0,986), P30 vs P60 (p= 0,993).

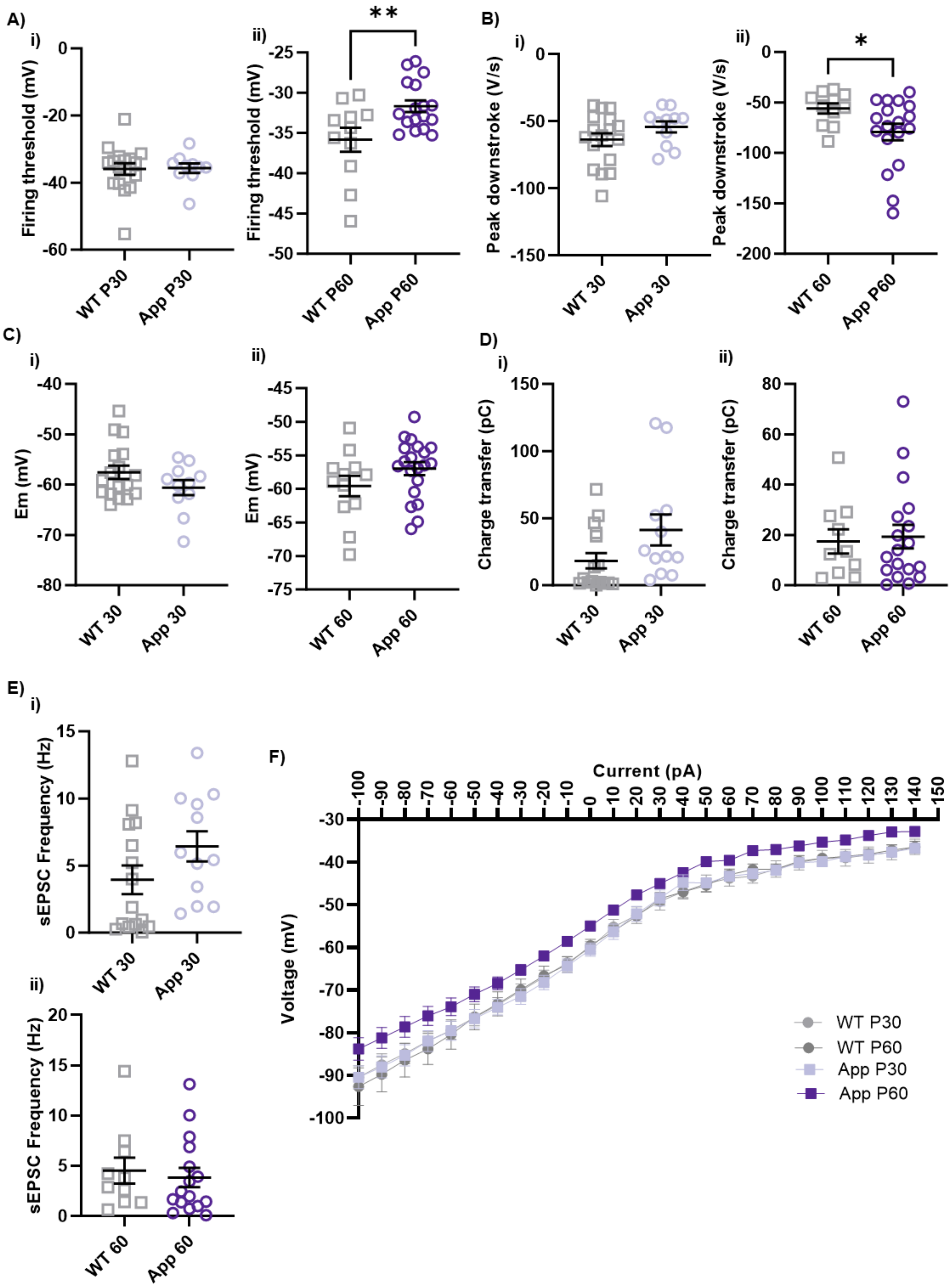
A)
i)
Firing threshold (mV)
0
-20
-40
-60
WT P30
App P30
ii)
**
Firing threshold (mV)
-25
-30
-35
-40
-45
-50
WT P60
App P60
B)
i)
Peak downstroke (V/s)
0
-50
-100
-150
WT 30
App 30
ii)
*
Peak downstroke (V/s)
0
-50
-100
-150
-200
WT 60
App 60
C)
i)
Em (mV)
-40
-50
-60
-70
-80
WT 30
App 30
ii)
Em (mV)
-45
-50
-55
-60
-65
-70
-75
WT 60
App 60
D)
i)
Charge transfer (pC)
150
100
50
0
WT 30
App 30
ii)
Charge transfer (pC)
80
60
40
20
0
WT 60
App 60
E)
i)
sEPSC Frequency (Hz)
15
10
5
0
WT 30
App 30
ii)
sEPSC Frequency (Hz)
20
15
10
5
0
WT 60
App 60
F)
Current (pA)
-100 -90 -80 -70 -60 -50 -40 -30 -20 -10 0 10 20 30 40 50 60 70 80 90 100 110 120 130 140 150
Voltage (mV)
-30
-40
-50
-60
-70
-80
-90
-100
WT P30
WT P60
App P30
App P60

37

**Supplementary figure 2 - Fast-spiking interneuron intrinsic properties and postsynaptic input are preserved in App$^{NL\text{-}G\text{-}F}$ P30.**

**A-E)** Scatter plots of FSI's intrinsic properties and post-synaptic input in the App$^{NL\text{-}G\text{-}F}$ group compared to age-matched WT. **Ai-ii)** T-test of firing threshold **i)** at P30 (WT n=17 vs App$^{NL\text{-}G\text{-}F}$ n=10 at P30 p=0,920) and **ii)** at P60 (WT n=11 vs App$^{NL\text{-}G\text{-}F}$ n=17 at P60 p=0,009). **Bi-ii)** T-test of peak downstroke **i)** at P30 (WT n=17 vs App$^{NL\text{-}G\text{-}F}$ n=11 at P30 p=0,180) and **ii)** at P60 (WT n=11 vs App$^{NL\text{-}G\text{-}F}$ n=18 at P60 p=0,046). **Ci-ii)** T-test of resting membrane potential (Em0) **i)** at P30 (WT n=17 vs App$^{NL\text{-}G\text{-}F}$ n=11 at P30 p=0,149) and **ii)** at P60 (WT n=12 vs App$^{NL\text{-}G\text{-}F}$ n=17 at P60 p=0,144). **Di-ii)** T-test of charge transfer **i)** at P30 (WT n=16 vs App$^{NL\text{-}G\text{-}F}$ n=12 at P30 p=0,065) and **ii)** at P60 (WT n=10 vs App$^{NL\text{-}G\text{-}F}$ n=18 at P60 p=0,800). **Ei-ii)** T-test of sEPSC frequency **i)** at P30 (WT n=16 vs App$^{NL\text{-}G\text{-}F}$ n=12 at P30 p=0,132) and **ii)** at P60 (WT n=10 vs App$^{NL\text{-}G\text{-}F}$ n=18 p=0,664). **F)** Current-voltage plot showing the membrane response to each current step in FSI from WT P30, WT P60, App$^{NL\text{-}G\text{-}F}$ P30 and App$^{NL\text{-}G\text{-}F}$ P60. FSI from App$^{NL\text{-}G\text{-}F}$ P60 are more depolirized and follow a different curve if compared to WT P30 and P60 and App$^{NL\text{-}G\text{-}F}$ P30. Data are presented as mean ± SEM, "n" indicates the number of neurons. *p < 0,05, **p < 0,01, ***p < 0,001.

**A)**

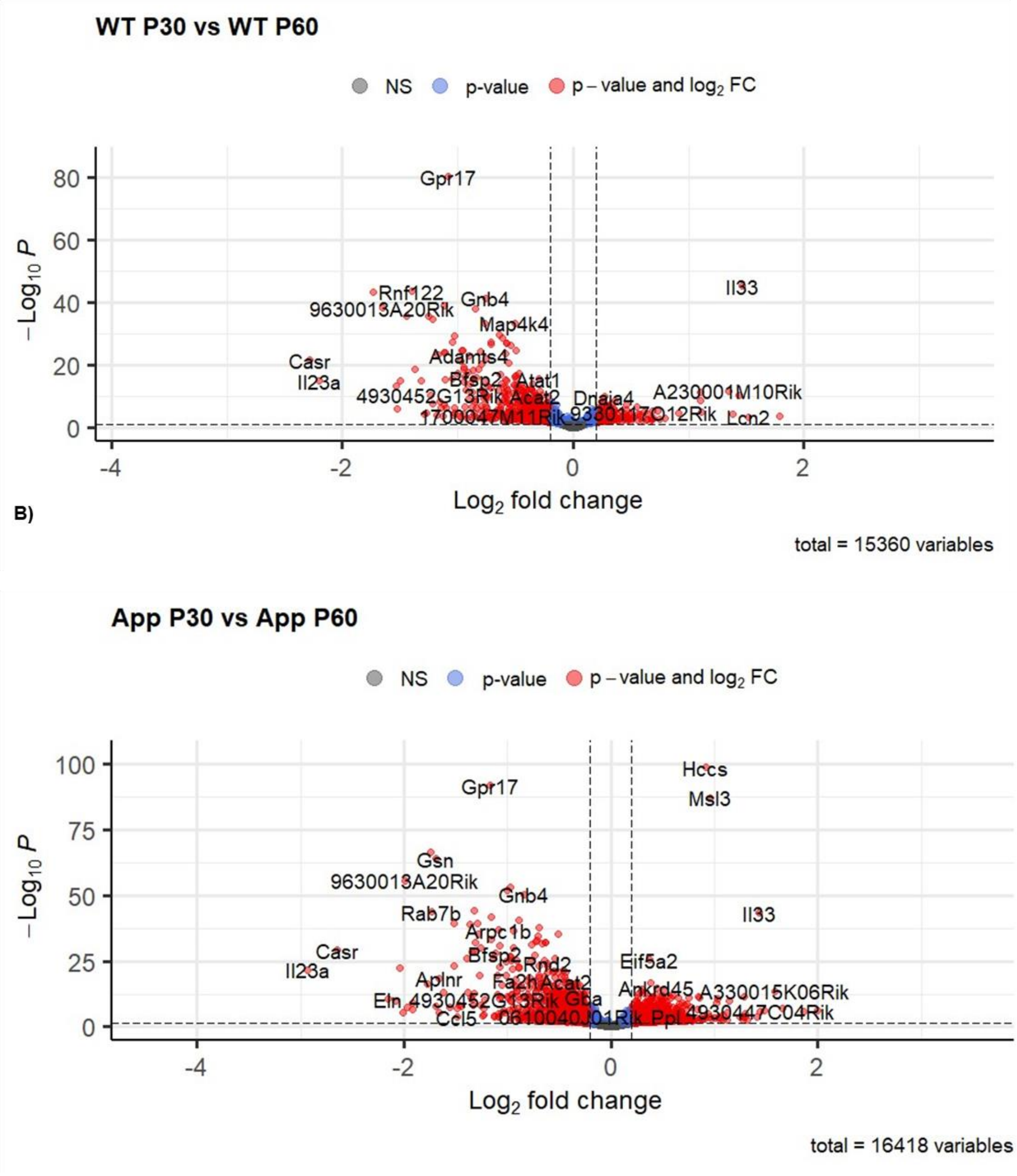


**Supplementary figure 3 – The bulk RNA sequencing analysis provides information on physiological hippocampal maturation and amyloid pathology progression.**

**A-B)** Volcano plots showing transcriptional differences due to **A)** the physiological maturation of the hippocampus (WT P30 n=4 vs WT P60 n=4; 998 DEG), **B)** the progression of amyloid pathology (App$^{NL\text{-}G\text{-}F}$ P30 n=4 vs App$^{NL\text{-}G\text{-}F}$ P60 n=4; 4246 DEG). Transcriptional differences were evaluated through a Wald test followed by apelgm shrinkage method and Benjamini & Hochberg multiple testing correction.

A)

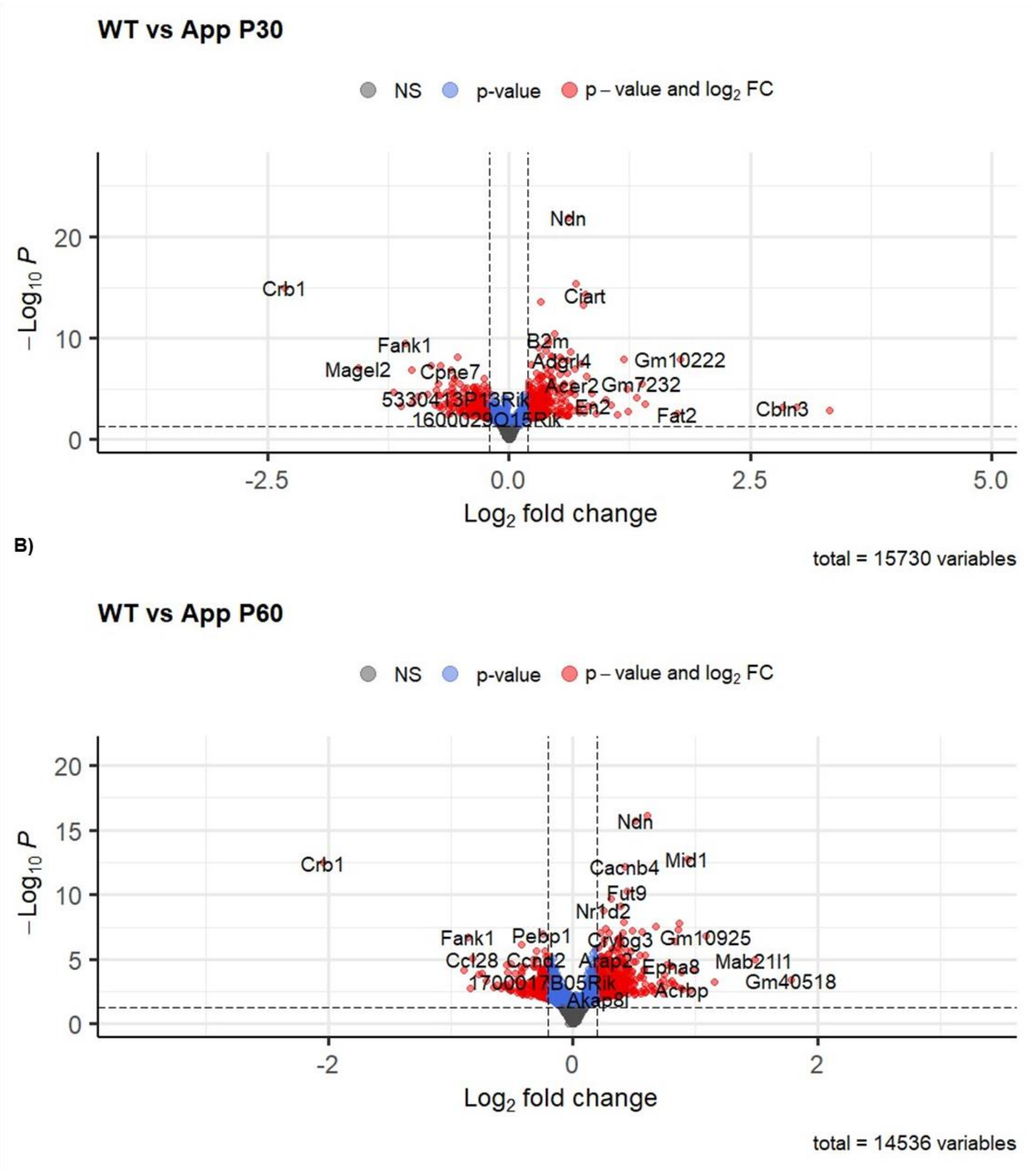

**Supplementary figure 4 – Bulk RNA sequencing analysis provides information on the effects of the $App^{NL\text{-}G\text{-}F}$ mutation at different ages.**

**A-B)** Volcano plots showing transcriptional differences due to the $App^{NL\text{-}G\text{-}F}$ mutation **A)** at P30 (WT P30 n=4 vs $App^{NL\text{-}G\text{-}F}$ P30 n=4; 510 DEG), and **B)** at P60 (WT P60 n=4 vs $App^{NL\text{-}G\text{-}F}$ P60 n=4; 930 DEG).Transcriptional differences were evaluated through a Wald test followed by apelgm shrinkage method and Benjamini & Hochberg multiple testing correction.

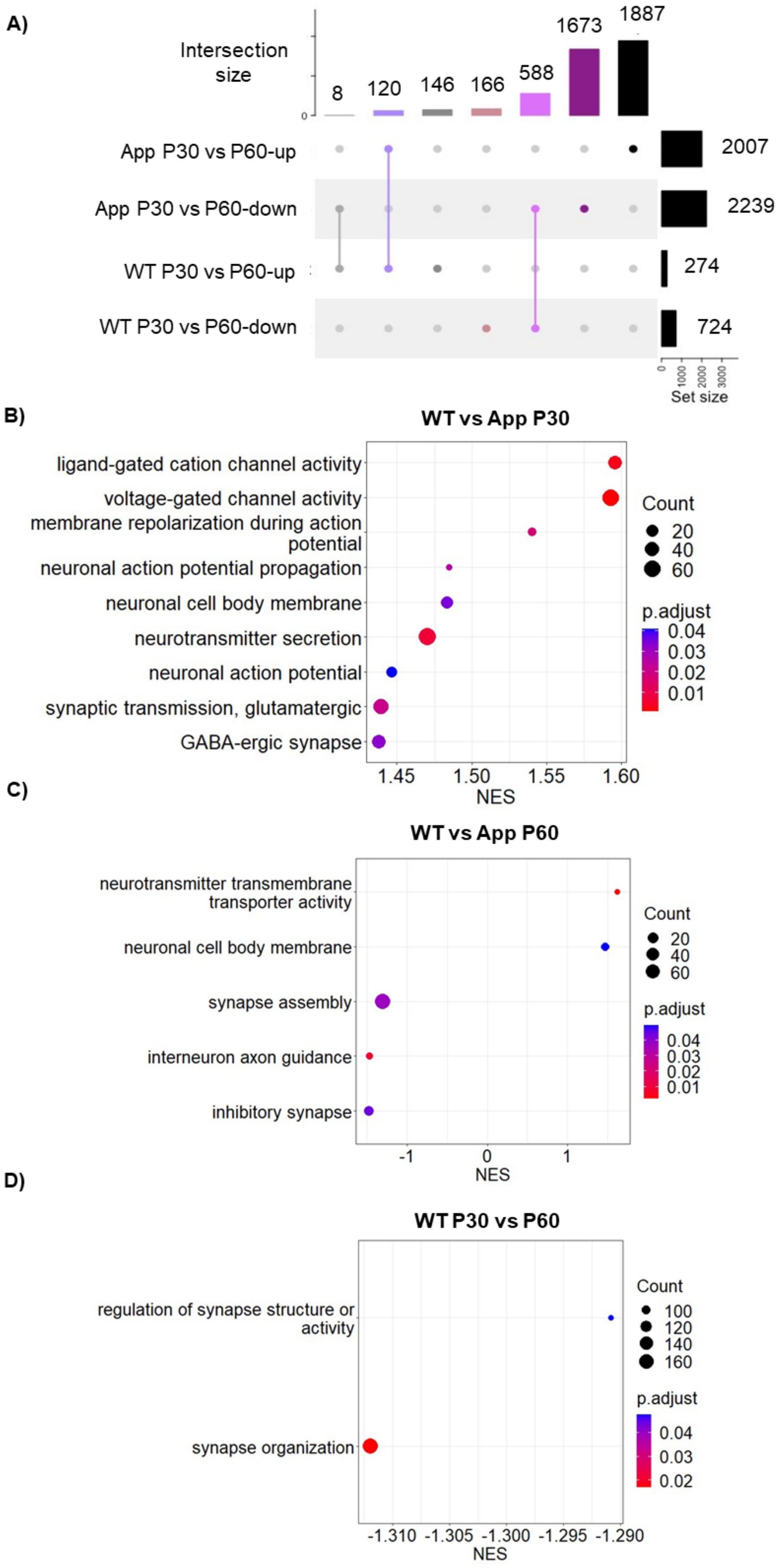

A)
Intersection size
8
120
146
166
588
1673
1887
App P30 vs P60-up
App P30 vs P60-down
WT P30 vs P60-up
WT P30 vs P60-down
2007
2239
274
724
0
1000
2000
3000
Set size
B)
WT vs App P30
ligand-gated cation channel activity
voltage-gated channel activity
membrane repolarization during action potential
neuronal action potential propagation
neuronal cell body membrane
neurotransmitter secretion
neuronal action potential
synaptic transmission, glutamatergic
GABA-ergic synapse
1.45
1.50
1.55
1.60
NES
Count
20
40
60
p.adjust
0.04
0.03
0.02
0.01
C)
WT vs App P60
neurotransmitter transmembrane transporter activity
neuronal cell body membrane
synapse assembly
interneuron axon guidance
inhibitory synapse
-1
0
1
NES
Count
20
40
60
p.adjust
0.04
0.03
0.02
0.01
D)
WT P30 vs P60
regulation of synapse structure or activity
synapse organization
-1.310
-1.305
-1.300
-1.295
-1.290
NES
Count
100
120
140
160
p.adjust
0.04
0.03
0.02

**Supplementary figure 5 – Gene set enrichment analysis exploring synaptic pathway changes during normal development and at different stages of amyloidogenic progression.**

**A)** Upset plot showing the number of upregulated and downregulated DEG in App$^{NL-G-F}$ P30 vs P60 and WT P30 vs P60 with respective number overlapping DEG (Intersection size). **B-D)** Enrichment plots of synaptic pathways in **A)** WT P30 (n=4) vs App$^{NL-G-F}$ P30 (n=4), **B)** WT P60 (n=4) vs App$^{NL-G-F}$ P60 (n=4) and **C)** WT P630 (n=4) vs WT P30 (n=4).

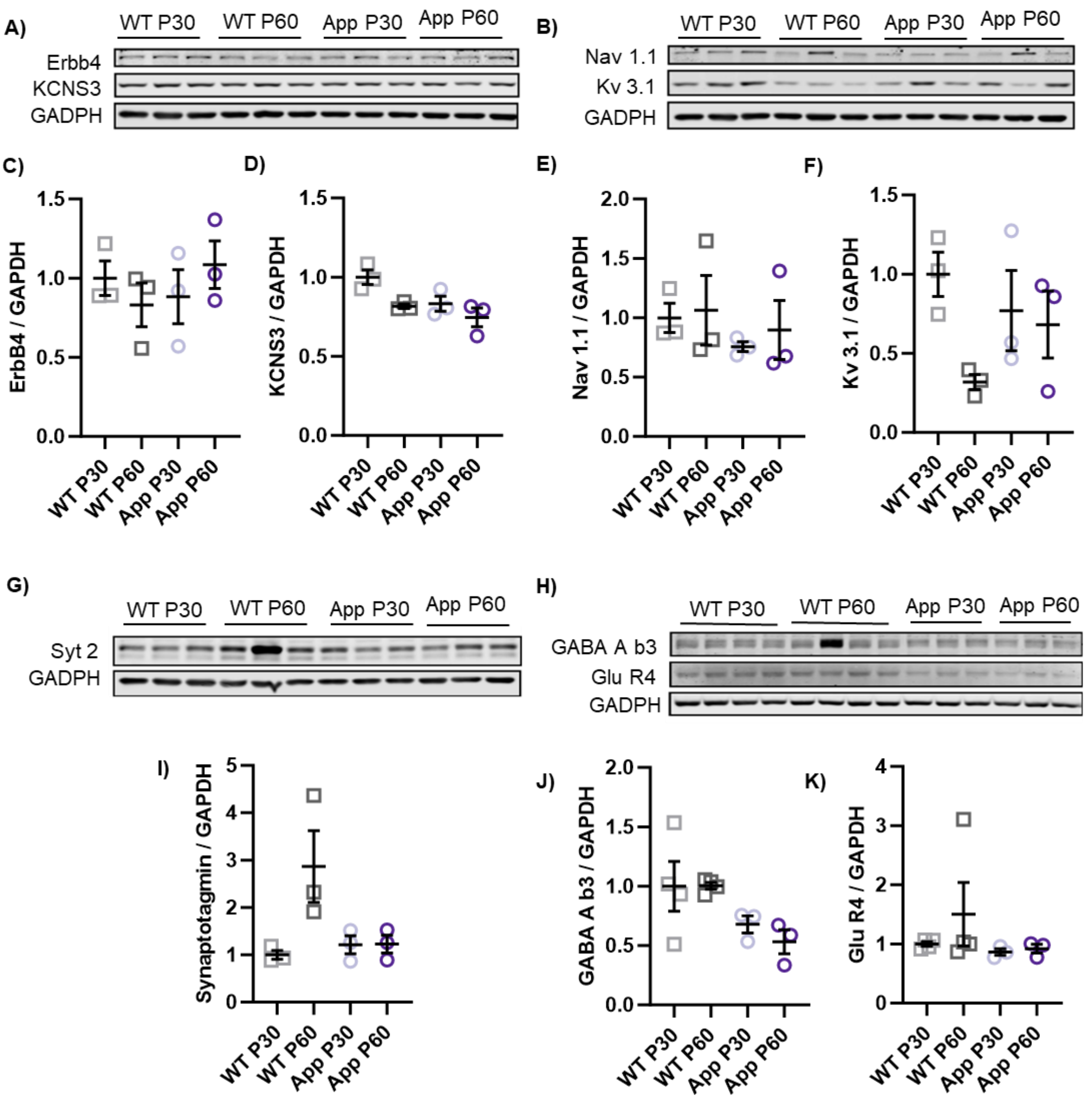


**Supplementary figure 6 – Synaptic and FSI related protein levels evaluated through western blot.**

**A, B, G, H)** Western blot membranes of **A)** Erbb4, NCNS3 and GADPH, **B)** Nav1.1, Kv3.1 and GADPH, **G)** Syt2 and GADPH and **H)** GABA A b3, Glu R4 and GADPH from **A, B, G**) WT P30 (n=3), WT P60 (n=3), $App^{NL\text{-}G\text{-}F}$ P30 (n=3), and $App^{NL\text{-}G\text{-}F}$ P60 (n=3) and **H)** WT P30 (n=4), WT P60 (n=4), $App^{NL\text{-}G\text{-}F}$ P30 (n=3), and $App^{NL\text{-}G\text{-}F}$ P60 (n=3). **C)** Scatter plot of Erbb4 normalized protein level showing WT P30 vs $App^{NL\text{-}G\text{-}F}$ P30 (p=0,927), WT P60 vs App$^{NL-G-F}$ P60 (p=0,818), App$^{NL-G-F}$ P30 vs App$^{NL-G-F}$ P60 (p=0,883). **D)** Scatter plot of KCNS3 normalized protein level showing WT P30 vs App$^{NL-G-F}$ P30 (p=0,086), WT P60 vs App$^{NL-G-F}$ P60 (p=0380), App$^{NL-G-F}$ P30 vs App$^{NL-G-F}$ P60 (p=0,380). **E)** Scatter plot of Nav1.1 normalized protein level showing WT P30 vs App$^{NL-G-F}$ P30 (p=0,888), WT P60 vs App$^{NL-G-F}$ P60 (p=0,924), App$^{NL-G-F}$ P30 vs App$^{NL-G-F}$ P60 (p=0,924). **F)** Scatter plot of Kv3.1 normalized protein level showing WT P30 vs App$^{NL-G-F}$ P30 (p=0,635), WT P60 vs App$^{NL-G-F}$ P60 (p=0,475), App$^{NL-G-F}$ P30 vs App$^{NL-G-F}$ P60 (p=0,740). **I)** Scatter plot of Syt2 normalized protein level showing WT P30 vs App$^{NL-G-F}$ P30 (p=0,920), WT P60 vs App$^{NL-G-F}$ P60 (p=0,062), App$^{NL-G-F}$ P30 vs App$^{NL-G-F}$ P60 (p=0,984). **J)** Scatter plot of GABA A b3 normalized protein level showing WT P30 vs App$^{NL-G-F}$ P30 (p=0,387), WT P60 vs App$^{NL-G-F}$ P60 (p=0,129), App$^{NL-G-F}$ P30 vs App$^{NL-G-F}$ P60 (p=0,890). **K)** Scatter plot of GluR4 normalized protein level showing WT P30 vs App$^{NL-G-F}$ P30 (p=0,990), WT P60 vs App$^{NL-G-F}$ P60 (p=0,592), App$^{NL-G-F}$ P30 vs App$^{NL-G-F}$ P60 (p=0,999). All scatter plots show statistics from one-way ANOVA with Holm–Sidak's multiple comparisons test. Data are presented as mean ± SEM, "n" indicates the number of animals.

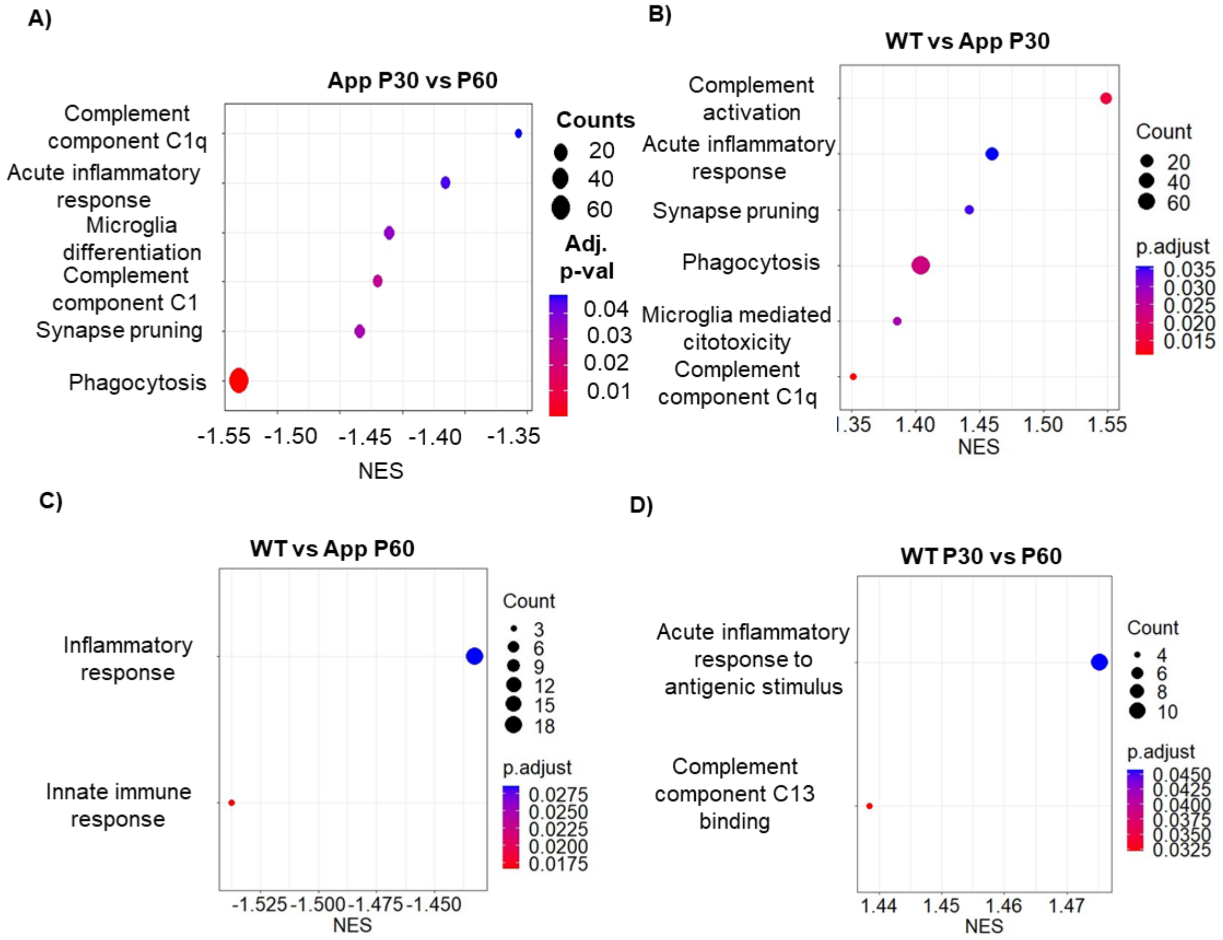


**Supplementary figure 7 – Gene set enrichment analysis exploring microglia- and inflammation-related pathway changes during normal development and at different stages of amyloidogenic progression.**

**A-D)** Enrichment plots of microglia- and inflammation-related pathways in **A)** App$^{NL-G-F}$ P30 (n=4) vs P60 (n=4), **B)** WT P30 (n=4) vs App$^{NL-G-F}$ P30 (n=4), **C)** WT P60 (n=4) vs App$^{NL-G-F}$ P60 (n=4), and **D** WT P30 (n=4) vs P60 (n=4).

A)

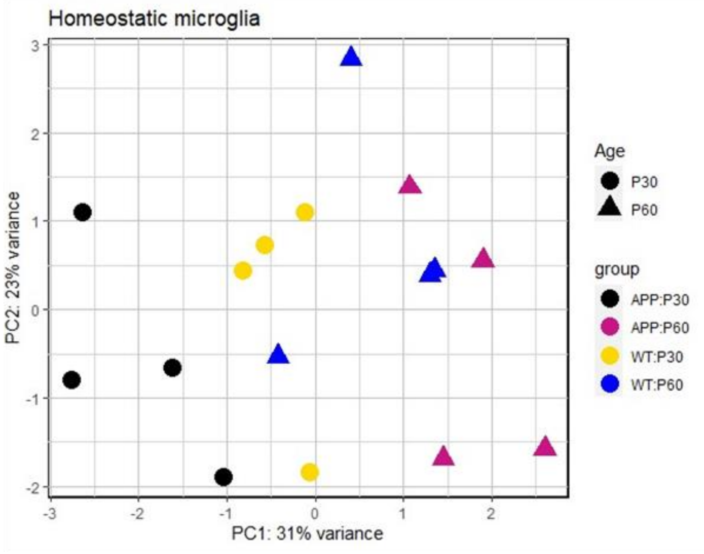
Homeostatic microglia
PC2: 23% variance
PC1: 31% variance
Age
P30
P60
group
APP:P30
APP:P60
WT:P30
WT:P60

B)

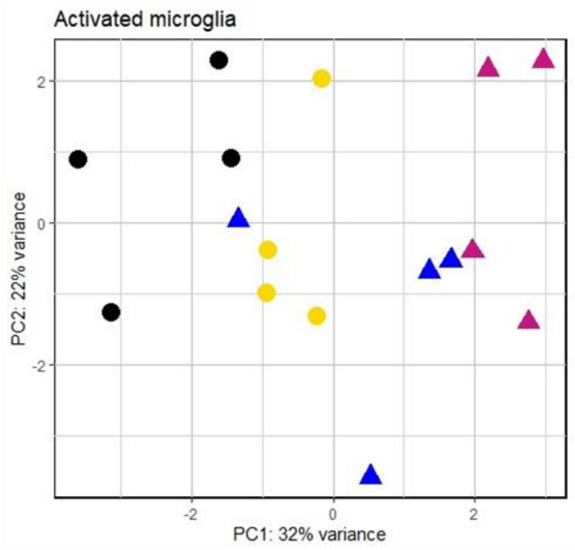
Activated microglia
PC2: 22% variance
PC1: 32% variance

C)

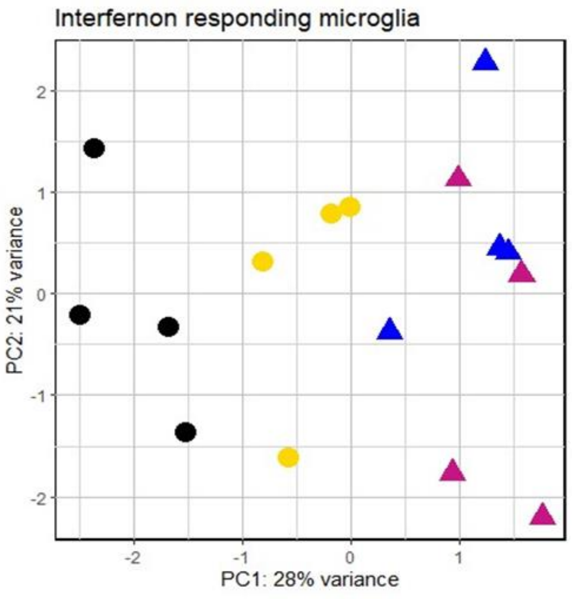
Interfernon responding microglia
PC2: 21% variance
PC1: 28% variance

**Supplementary figure 8 – Homeostatic, activated and interferon responding microglia profiles are different between pre-plaque and plaque stage of amyloid pathology progression.**

**A-C)** PCA for microglia profile representative genes including **A)** 751 genes for homeostatic and pruning microglia, **B)** 885 genes for activated microglia and **C)** 418 genes for interferon-responding microglia.

#### Videos

**Supplementary video 1 – 3D PCA plot of FSI intrinsic properties**

#### Tables

| Gene | Protein | Function | Log2 fold change | |
|---|---|---|---|---|
| | | | App P30vsP60 | WTvsAppP60 |
| Kcnc1 | Kv3.1 | Potassium channel with high activation threshold, and fast activation/deactivation kinetics. It is involved in the rapid repolarization of the action potential (Labro et al., 2015), and its presence in neurons correlates with high-frequency firing (Kaczmarek & Zhang, 2017) like FSI. Kv3.1 is highly expressed in GI in the hippocampus. the proper functionality of Kv3.1 has been connected to gamma oscillation restoration in mouse hippocampal slices after acute Ab exposure (Andrade-Talavera et al., 2020). | 0,282435212<br>p=0,0002<br>*** | 0,145549249<br>p= 0,07073608 |
| Kcns3 | Kv9.3 | Potassium channel subunit that closely correlates with FSI's firing feature (Georgiev et al., 2012). Kcns3 deficiency has been shown to impair the recruitment of FSI during gamma oscillations in the cortex (Miyamae et al., 2021) | 0,216522711<br>p=0,031101797<br>* | 0,197639332<br>p=0,061690371 |
| Scn1a | Nav1.1 | Sodium channel connected to fast axonal propagation features in FSI (Martin et al., 2010). Mutations of Scn1a have been shown to alter brain oscillations in the gamma band in clinical studies (Holmes et al., 2012) | 0,393441525<br>p=0,00000193<br>**** | 0,36237221<br>p=0,000131862<br>*** |
| Erbb4 | Erbb4 | Tyrosine kinase receptor expressed selectively by FSI in the hippocampus (Vullhorst et al., 2009). Its activation increases GABA release (Robinson et al., | 0,269006844<br>p=0,016316214 | 0,211166543<br>p=0,066469927 |

| | | | | |
|---|---|---|---|---|
| | | 2022) and its activity has been connected to gamma oscillation regulation in a physiological state (Tan et al., 2018) in AD models (Zhang et al., 2017) | * | |
| GABRA1 | GABA A receptor a1 | GABRA1 encodes one of the most abundant subunits forming the GABA A receptor, namely alpha1 (Olsen & Sieghart, 2009), which has been connected to gamma oscillations onset (Kujuala, 2015). | 0,25191162<br>p=0,001699343<br>** | 0,21223797<br>p=0,023832647<br>* |
| Syt2 | Syt2 | Synaptotagmin2 belongs to the calcium-binding subgroup of the synaptotagmin family, a group of presynaptic proteins that plays a fundamental role in neurotransmitter exocytosis (Maximov, 2009). Syt2 expression is associated with fast, synchronized vesicle release (Chen et al., 2017; Pang et al., 2006). and it has been proven to be a reliable marker for FSI in multiple brain areas (Chen et al., 2017; Sommeijer & Levelt, 2012). | 0,209824837<br>p=0,118514546 | 0,864058576<br>p= 0,0000153<br>**** |

**Supp. Table 1 – Synaptic and FSI related genes**

Genes involved in synaptic activity and FSI's functionality. Transcriptional differences were evaluated through a Wald test followed by apelgm shrinkage method and Benjamini & Hochberg multiple testing correction. *p < 0,05, **p < 0,01, ***p < 0,001, **** p < 0,0001.

**Supp. Table 2 – Gene-set enrichment analysis**

Table showing the significantly enriched pathways for all comparisons.

**Supp. Table 3 – Microglia signature genes**

Table of 2054 microglia expressed genes with respective raw counts extracted from the bulk RNA sequencing dataset. Sheets listing 751 genes for homeostatic and pruning microglia, 885 genes for activated microglia, and 418 genes for interferon-related microglia and respective PCA coordinates are included.

#### Supplementary tables of graph data

| | Source of variation | Test | P value |
|---|---|---|---|
| **Two-way ANOVA test** | Row factor | F (3, 49) = 4,751 | P=0,0055 |
| | Column factor | F (1, 49) =15,65 | P=0,0002 |

| **Fig 1 Panel C** | Gamma power | | | | | |
|---|---|---|---|---|---|---|
| | WT | | | $App^{NL-G-F}$ | | |
| Age | Mean | SD | n | Mean | SD | n |
| P30 | 2,768 | 0,952 | 6 | 2,175 | 0,708 | 6 |
| P40 | 2,512 | 1,330 | 6 | 1,527 | 0,300 | 6 |
| P50 | 2,105 | 0,860 | 6 | 1,249 | 0,619 | 9 |
| P60 | 2,027 | 0,629 | 7 | 1,180 | 0,521 | 11 |

| **Holm-Sidak's multiple comparisons test** | | |
|---|---|---|
| Comparison | Significant | P value |
| WT vs App P30 | No | 0,124 |
| WT vs App P40 | No | 0,172 |
| WT vs App P50 | Yes | 0,017 |
| WT vs App P60 | Yes | 0,010 |
| WT P30 vs P40 | No | 0,286 |
| WT P30 vs P50 | No | 0,304 |
| WT P30 vs P60 | No | 0,255 |
| App P30 vs P40 | No | 0,326 |
| App P30 vs P50 | Yes | 0,049 |
| App P30 vs P60 | Yes | 0,026 |

| | Source of variation | Test | P value |
|---|---|---|---|
| Two-way ANOVA test | Column factor | F (1, 16) = 5,761 | P=0,0289 |

| **Fig 1 Panel D** | c-Fos normalized expression | | | | | | | |
|---|---|---|---|---|---|---|---|---|
| | Control | | | Kainic Acid | | | Holm-Sidak's multiple comparisons test | |
| Group | Mean | SD | n | Mean | SD | n | Significant | P value |
| WT | 1 | 0,219 | 4 | 1,817 | 0,107 | 4 | Yes | 0,0035 |
| $App^{NL-G-F}$ P30 | 0,998 | 0,297 | 3 | 0,880 | 0,213 | 3 | No | 0,9990 |
| $App^{NL-G-F}$ P60 | 0,997 | 0,208 | 4 | 1,063 | 0,366 | 4 | No | 0,9994 |

| | Source of variation | Test | P value |
|---|---|---|---|
| Two-way ANOVA test | Column factor | F (1, 16) = 6,287 | P=0,0233 |

| **Fig 1 Panel E** | NPAS4 normalized expression | | | | | | | |
|---|---|---|---|---|---|---|---|---|
| | Control | | | Kainic Acid | | | Holm-Sidak's multiple comparisons test | |
| Group | Mean | SD | n | Mean | SD | n | Significant | P value |
| WT | 0,999 | 0,234 | 4 | 2,084 | 0,309 | 4 | Yes | <0,0001 |
| $App^{NL-G-F}$ P30 | 1 | 0,136 | 3 | 0,849 | 0,106 | 3 | No | 0,9827 |
| $App^{NL-G-F}$ P60 | 1 | 0,224 | 4 | 0,833 | 0,280 | 4 | No | 0,9827 |

| **Fig 2 Panel D-H** | FSI intrinsic properties and post synaptic input | | | | | | | | | |
|---|---|---|---|---|---|---|---|---|---|---|
| | $App^{NL-G-F}$ P30 | | | $App^{NL-G-F}$ P60 | | | | | | |
| | Mean | SD | n | Mean | SD | n | Test | Significant | P value | Difference ± SEM |
| Resting membrane potential (mV) | -60,60 | 4,912 | 11 | -56,99 | 4,335 | 17 | Unpaired T-test | Yes | 0,0291 | 3,708 ± 1,622 |
| Firing threshold (mV) | -35,67 | 4,545 | 11 | -31,67 | 3,021 | 17 | Unpaired T-test | Yes | 0,0109 | 3,995 ± 1,452 |
| Peak downstroke (V/s) | -55,77 | 13,27 | 11 | -80,8 | 53,43 | 17 | Unpaired T-test | Yes | 0,0488 | -24,30 ± 11,73 |
| Charge transfer (pC) | 40,66 | 40,03 | 11 | 19,33 | 19,82 | 18 | Mann-Whitney | yes | 0,0275 | -13,39 |
| sEPSC frequency (Hz) | 6,889 | 3,752 | 11 | 3,821 | 3,809 | 18 | Mann-Whitney | yes | 0,0220 | -3,208 |

| **Fig 3 Panel F** | GABA R1 alpha normalized expression | | | One-way ANOVA test | F= 10,12 | P= 0,0023 |
|---|---|---|---|---|---|---|
| | | | | Holm-Sidak's multiple comparisons test | | |
| Group | Mean | SD | n | Comparison | Significant | P value |
| WT P30 | 1 | 0,303 | 4 | WT P30 vs App P30 | Yes | 0,032 |
| WT P60 | 1,158 | 0,305 | 4 | WT P60 vs App P60 | Yes | 0,007 |
| $App^{NL-G-F}$ P30 | 0,399 | 0,194 | 3 | WT P30 vs P60 | No | 0,635 |
| $App^{NL-G-F}$ P60 | 0,288 | 0,035 | 3 | App P30 vs P60 | No | 0,635 |

| **Fig 3 Panel G** | Glu A1 normalized expression | | | One-way ANOVA test | F= 13,36 | P= 0,0008 |
|---|---|---|---|---|---|---|
| | | | | Holm-Sidak's multiple comparisons test | | |
| Group | Mean | SD | n | Comparison | Significant | P value |
| WT P30 | 1 | 0,201 | 4 | WT P30 vs App P30 | Yes | 0,009 |
| WT P60 | 0,893 | 0,144 | 4 | WT P60 vs App P60 | Yes | 0,005 |
| $App^{NL-G-F}$ P30 | 0,553 | 0,095 | 3 | WT P30 vs P60 | No | 0,376 |
| $App^{NL-G-F}$ P60 | 0,397 | 0,036 | 3 | App P30 vs P60 | No | 0,376 |

| **Fig 4 Panel C** | Iba1 normalized expression | | | One-way ANOVA test | F= 9,081 | P= 0,0008 |
|---|---|---|---|---|---|---|
| | | | | Holm-Sidak's multiple comparisons test | | |
| Group | Mean | SD | n | Comparison | Significant | P value |
| WT P30 | 0,787 | 0,139 | 5 | WT P30 vs App P30 | Yes | 0,009 |
| WT P60 | 0,607 | 0,193 | 5 | WT P60 vs App P60 | No | 0,282 |
| $App^{NL-G-F}$ P30 | 1,290 | 0,237 | 5 | WT P30 vs P60 | No | 0,368 |
| $App^{NL-G-F}$ P60 | 0,832 | 0,262 | 6 | App P30 vs P60 | Yes | 0,011 |

| **Fig 5 Panel K** | Microglia density (cell/nm$^2$) | | | One-way ANOVA test | F= 9,389 | P= 0,0001 |
|---|---|---|---|---|---|---|
| | | | | Holm-Sidak's multiple comparisons test | | |
| Group | Mean | SD | n | Comparison | Significant | P value |
| WT P30 | 265,9 | 52,96 | 11 | WT P30 vs App P30 | Yes | 0,036 |
| WT P60 | 240,7 | 65,23 | 9 | WT P60 vs App P60 | No | 0,096 |
| App$^{NL-G-F}$ P30 | 322,8 | 22,59 | 6 | WT P30 vs P60 | No | 0,253 |
| App$^{NL-G-F}$ P60 | 192,6 | 32,09 | 11 | App P30 vs P60 | Yes | <0,0001 |

| **Fig 5 Panel L** | Microglia-FSI body contact area (nm$^2$) | | | | | |
|---|---|---|---|---|---|---|
| | | | | Unpaired t-test (one tailed) | | |
| Group | Mean | SD | N | Comparison | Significant | P value |
| WT P30 | 12,69 | 11,52 | 6 | WT P30 vs App P30 | No | 0,036 |
| WT P60 | 9,06 | 9,23 | 6 | WT P60 vs App P60 | Yes | 0,039 |
| App$^{NL-G-F}$ P30 | 8,67 | 8,92 | 6 | WT P30 vs P60 | No | 0,280 |
| App$^{NL-G-F}$ P60 | 1,53 | 1,79 | 6 | App P30 vs P60 | Yes | 0,041 |

| **Fig 5 Panel M** | Pearson r correlation test (Microglia – plaques in App$^{NL-G-F}$ P60) | | | | | | |
|---|---|---|---|---|---|---|---|
| | Mean | SD | n | Pair | Pearson r | Significant | P value |
| Microglia density (cell/nm$^2$) | 202,2 | 26,76 | 9 | Microglia density – Microglia density | 1 | / | / |
| Plaque density (n/nm$^2$) | 14,66 | 4,081 | 9 | Microglia density - Plaque density | -0,290 | No | 0,449 |
| Normalized plaque area | 0,574 | 0,141 | 9 | Microglia density - Normalized plaque area | -0,671 | Yes | 0,048 |

| **Fig 5 Panel N** | Pearson r correlation test (Microglia-FSI body contact area – FIS intrinsic properties) | | | | | | |
|---|---|---|---|---|---|---|---|
| | Mean | SD | n | Pair | Pearson r | Significant | P value |
| Microglia-FSI body contact area (nm$^2$) | 8,022 | 8,331 | 19 | Microglia-FSI body contact area – Microglia-FSI body contact area | 1 | / | / |
| Firing threshold (mV) | -34,73 | 4,200 | 19 | Microglia-FSI body contact area - Firing threshold | -0,548 | Yes | 0,015 |

| | | | | | | | |
|---|---|---|---|---|---|---|---|
| Resting membrane potential (mV) | -56,92 | 4,614 | 19 | Microglia-FSI body contact area - Resting membrane potential | -0,188 | No | 0,456 |
| Peak downstroke (V/s) | -76,70 | 32,78 | 19 | Microglia-FSI body contact area - Peak downstroke | -0,023 | No | 0,927 |
| N of spikes | 281,9 | 148,0 | 19 | Microglia-FSI body contact area - N of spikes | 0,519 | Yes | 0,023 |

| | Source of variation | Test | P value |
|---|---|---|---|
| **Two-way ANOVA test** | Row factor | F (3, 49) = 0,859 | P=0,4687 |
| | Column factor | F (3, 49) =3,822 | P=0,0563 |

| **Supp Fig 1 Panel A** | Frequency variance (Hz) | | | | | | **Holm-Sidak's multiple comparisons test** | | |
|---|---|---|---|---|---|---|---|---|---|
| | WT | | | $App^{NL-G-F}$ | | | | | |
| Age | Mean | SD | n | Mean | SD | n | Comparison | Significant | P value |
| P30 | 8,413 | 2,068 | 6 | 11,580 | 3,877 | 6 | WT vs App P30 | No | 0,831 |
| P40 | 14,745 | 3,703 | 6 | 8,628 | 2,358 | 6 | WT vs App P40 | No | 0,161 |
| P50 | 12,315 | 3,838 | 6 | 10,363 | 3,350 | 9 | WT vs App P50 | No | 0,981 |
| P60 | 13,301 | 3,887 | 7 | 10,541 | 2,739 | 11 | WT vs App P60 | No | 0,802 |
| | | | | | | | WT P30 vs P40 | No | 0,125 |
| | | | | | | | WT P30 vs P50 | No | 0,630 |
| | | | | | | | WT P30 vs P60 | No | 0,202 |
| | | | | | | | App P30 vs P40 | No | 0,881 |
| | | | | | | | App P30 vs P50 | No | 0,989 |
| | | | | | | | App P30 vs P60 | No | 0,989 |

| | Source of variation | Test | P value |
|---|---|---|---|
| **Two-way ANOVA test** | Row factor | F (3, 49) = 3,156 | P=0,0329 |
| | Column factor | F (1, 49) =0,02149 | P=0,8841 |

| **Supp Fig 1 Panel B** | Peak frequency (Hz) | | | | | | **Holm-Sidak's multiple comparisons test** | | |
|---|---|---|---|---|---|---|---|---|---|
| | WT | | | $App^{NL-G-F}$ | | | | | |
| Age | Mean | SD | n | Mean | SD | n | Comparison | Significant | P value |
| P30 | 25,838 | 1,917 | 6 | 27,178 | 2,930 | 6 | WT vs App P30 | No | 0,994 |
| P40 | 28,415 | 3,484 | 6 | 26,720 | 2,267 | 6 | WT vs App P40 | No | 0,993 |
| P50 | 30,484 | 3,699 | 6 | 28,992 | 2,303 | 9 | WT vs App P50 | No | 0,993 |
| P60 | 27,291 | 2,431 | 7 | 28,699 | 2,595 | 11 | WT vs App P60 | No | 0,993 |
| | | | | | | | WT P30 vs P40 | No | 0,928 |
| | | | | | | | WT P30 vs P50 | No | 0,141 |

| | | |
|---|---|---|
| WT P30 vs P60 | No | 0,993 |
| App P30 vs P40 | No | 0,992 |
| App P30 vs P50 | No | 0,986 |
| App P30 vs P60 | No | 0,993 |

| **Supp Fig 2 Panel A** | Firing threshold (mV) | | | | | | | | | |
|---|---|---|---|---|---|---|---|---|---|---|
| | WT | | | App$^{NL-G-F}$ | | | Unpaired T-test | | | |
| | Mean | SD | n | Mean | SD | n | Comparison | Significant | P value | Difference ± SEM |
| P30 | -35,92 | 7,112 | 17 | -35,67 | 4,545 | 10 | WT vs App P30 | No | 0,9198 | 0,2556 ± 2,515 |
| P60 | -35,84 | 4,925 | 11 | -31,67 | 3,021 | 17 | WT vs App P60 | Yes | 0,0099 | 4,162 ± 1,452 |

| **Supp Fig 2 Panel B** | Peak downstroke (V/s) | | | | | | | | | |
|---|---|---|---|---|---|---|---|---|---|---|
| | WT | | | App$^{NL-G-F}$ | | | Unpaired T-test | | | |
| | Mean | SD | n | Mean | SD | n | Comparison | Significant | P value | Difference ± SEM |
| P30 | -63,60 | 19,90 | 17 | -54,12 | 13,74 | 11 | WT vs App P30 | No | 0,1798 | 9,487 ± 6,882 |
| P60 | -55,89 | 16,49 | 11 | -79,28 | 34,54 | 18 | WT vs App P60 | Yes | 0,0458 | -23,39± 11,17 |

| **Supp Fig 2 Panel C** | Resting membrane potential (mV) | | | | | | | | | |
|---|---|---|---|---|---|---|---|---|---|---|
| | WT | | | App$^{NL-G-F}$ | | | Unpaired T-test | | | |
| | Mean | SD | n | Mean | SD | n | Comparison | Significant | P value | Difference ± SEM |
| P30 | -57,57 | 5,456 | 17 | -60,60 | 4,912 | 11 | WT vs App P30 | No | 0,1487 | -3,026 ± 2,033 |
| P60 | -59,57 | 5,251 | 12 | -56,99 | 4,355 | 17 | WT vs App P60 | No | 0,1444 | 2,574 ± 1,717 |

| **Supp Fig 2 Panel D** | Charge transfer (pC) | | | | | | | | | |
|---|---|---|---|---|---|---|---|---|---|---|
| | WT | | | $App^{NL-G-F}$ | | | Unpaired T-test | | | |
| | Mean | SD | n | Mean | SD | n | Comparison | Significant | P value | Difference ± SEM |
| P30 | 18,25 | 23,02 | 16 | 41,28 | 39,93 | 12 | WT vs App P30 | No | 0,0651 | 23,03 ± 11,96 |
| P60 | 17,48 | 15,16 | 10 | 19,33 | 19,82 | 18 | WT vs App P60 | No | 0,800 | 1,851 ± 7,232 |

| **Supp Fig 2 Panel D** | sEPSC frequency (Hz) | | | | | | | | | |
|---|---|---|---|---|---|---|---|---|---|---|
| | WT | | | $App^{NL-G-F}$ | | | Unpaired T-test | | | |
| | Mean | SD | n | Mean | SD | n | Comparison | Significant | P value | Difference ± SEM |
| P30 | 3,949 | 4,111 | 16 | 6,435 | 3,909 | 12 | WT vs App P30 | No | 0,1232 | 2,486 ± 1,558 |
| P60 | 4,523 | 4,111 | 10 | 3,821 | 3,586 | 18 | WT vs App P60 | No | 0,6413 | -0,7020 ± 1,489 |

| **Supp Fig 6 Panel C** | ErbB4 normalized expression | | | One-way ANOVA test | F= 0,633 | P= 0,6139 |
|---|---|---|---|---|---|---|
| | | | | Holm-Sidak's multiple comparisons test | | |
| Group | Mean | SD | n | Comparison | Significant | P value |
| WT P30 | 1 | 0,189 | 3 | WT P30 vs App P30 | No | 0,927 |
| WT P60 | 0,832 | 0,239 | 3 | WT P60 vs App P60 | No | 0,818 |
| $App^{NL-G-F}$ P30 | 0,883 | 0,295 | 3 | WT P30 vs P60 | No | 0,896 |
| $App^{NL-G-F}$ P60 | 1,085 | 0,260 | 3 | App P30 vs P60 | No | 0,883 |

| **Supp Fig 6 Panel D** | KCNS3 normalized expression | | | One-way ANOVA test | F= 5,754 | P= 0,0214 |
|---|---|---|---|---|---|---|
| | | | | Holm-Sidak's multiple comparisons test | | |
| Group | Mean | SD | n | Comparison | Significant | P value |
| WT P30 | 1 | 0,079 | 3 | WT P30 vs App P30 | No | 0,086 |
| WT P60 | 0,817 | 0,025 | 3 | WT P60 vs App P60 | No | 0,380 |
| $App^{NL-G-F}$ P30 | 0,832 | 0,082 | 3 | WT P30 vs P60 | No | 0,077 |
| $App^{NL-G-F}$ P60 | 0,747 | 0,102 | 3 | App P30 vs P60 | No | 0,380 |

| **Supp Fig 6 Panel E** | Nav1.1 normalized expression | One-way ANOVA test | F= 4,366 | P= 0,7329 |
|---|---|---|---|---|
| | | Holm-Sidak's multiple comparisons test | | |

| Group | Mean | SD | n | Comparison | Significant | P value |
|---|---|---|---|---|---|---|
| WT P30 | 1 | 0,214 | 3 | WT P30 vs App P30 | No | 0,888 |
| WT P60 | 1,065 | 0,507 | 3 | WT P60 vs App P60 | No | 0,924 |
| App$^{NL-G-F}$ P30 | 0,757 | 0,074 | 3 | WT P30 vs P60 | No | 0,924 |
| App$^{NL-G-F}$ P60 | 0,897 | 0,432 | 3 | App P30 vs P60 | No | 0,924 |

| **Supp Fig 6 Panel F** | Kv3.1 normalized expression | | | One-way ANOVA test | F= 2,445 | P= 0,1388 |
|---|---|---|---|---|---|---|
| | | | | Holm-Sidak's multiple comparisons test | | |
| Group | Mean | SD | n | Comparison | Significant | P value |
| WT P30 | 1 | 0,242 | 3 | WT P30 vs App P30 | No | 0,635 |
| WT P60 | 0,318 | 0,083 | 3 | WT P60 vs App P60 | No | 0,475 |
| App$^{NL-G-F}$ P30 | 0,770 | 0,440 | 3 | WT P30 vs P60 | No | 0,110 |
| App$^{NL-G-F}$ P60 | 0,682 | 0,367 | 3 | App P30 vs P60 | No | 0,740 |

| **Supp Fig 6 Panel I** | Syt2 normalized expression | | | One-way ANOVA test | F= 4,595 | P= 0,0376 |
|---|---|---|---|---|---|---|
| | | | | Holm-Sidak's multiple comparisons test | | |
| Group | Mean | SD | n | Comparison | Significant | P value |
| WT P30 | 1 | 0,161 | 3 | WT P30 vs App P30 | No | 0,920 |
| WT P60 | 2,868 | 1,314 | 3 | WT P60 vs App P60 | No | 0,062 |
| App$^{NL-G-F}$ P30 | 1,215 | 0,327 | 3 | WT P30 vs P60 | No | 0,050 |
| App$^{NL-G-F}$ P60 | 1,226 | 0,320 | 3 | App P30 vs P60 | No | 0,984 |

| **Supp Fig 6 Panel J** | GABA A b3 normalized expression | | | One-way ANOVA test | F= 3,002 | P= 0,0816 |
|---|---|---|---|---|---|---|
| | | | | Holm-Sidak's multiple comparisons test | | |
| Group | Mean | SD | n | Comparison | Significant | P value |
| WT P30 | 1 | 0,421 | 4 | WT P30 vs App P30 | No | 0,387 |
| WT P60 | 1,004 | 0,057 | 4 | WT P60 vs App P60 | No | 0,129 |
| App$^{NL-G-F}$ P30 | 0,679 | 0,126 | 3 | WT P30 vs P60 | No | >0,999 |
| App$^{NL-G-F}$ P60 | 0,532 | 0,532 | 3 | App P30 vs P60 | No | 0,890 |

| **Supp Fig 6 Panel K** | Glu R4 normalized expression | | | One-way ANOVA test | F= 0,9032 | P= 0,4734 |
|---|---|---|---|---|---|---|
| | | | | Holm-Sidak's multiple comparisons test | | |
| Group | Mean | SD | n | Comparison | Significant | P value |
| WT P30 | 1 | 0,078 | 4 | WT P30 vs App P30 | No | 0,990 |
| WT P60 | 1,503 | 1,071 | 4 | WT P60 vs App P60 | No | 0,592 |
| App$^{NL-G-F}$ P30 | 0,865 | 0,094 | 3 | WT P30 vs P60 | No | 0,640 |
| App$^{NL-G-F}$ P60 | 0,922 | 0,128 | 3 | App P30 vs P60 | No | 0,999 |
